# Nucleosome structural variations in interphase and metaphase chromosomes

**DOI:** 10.1101/2020.11.12.380386

**Authors:** Yasuhiro Arimura, Rochelle M. Shih, Ruby Froom, Hironori Funabiki

## Abstract

Structural heterogeneity of nucleosomes in functional chromosomes is unknown. Here we report cryo-EM structures of nucleosomes isolated from interphase and metaphase chromosomes at up to 3.4 Å resolution. Averaged chromosomal nucleosome structures are highly similar to canonical left-handed recombinant nucleosome crystal structures, with DNA being selectively stabilized at two defined locations. Compared to free mono-nucleosomes, which exhibit diverse linker DNA angles and large structural variations in H3 and H4, chromosomal nucleosome structures are much more uniform, characterized by a closed linker DNA angle with interactions between the H2A C-terminal tail and DNA. Exclusively for metaphase nucleosomes, structures of the linker histone H1.8 at the on-dyad position of nucleosomes can be reconstituted at 4.4 Å resolution. We also report diverse minor nucleosome structural variants with rearranged core histone configurations, which are more prevalent in metaphase than in interphase chromosomes. This study presents structural characteristics of nucleosomes in interphase and mitotic chromosomes.

**Highlights:**

- 3.4~ Å resolution nucleosome structures from interphase and metaphase chromosomes
- Nucleosome structures in chromosomes are more uniform than in free mono-nucleosomes
- Histone H1.8 binds to the nucleosome dyad axis in metaphase chromosomes
- Nucleosome structural variants are more prevalent in metaphase than in interphase

**NOTES TO READERS:** We would like to emphasize the importance of supplemental movies S1-S3, which should greatly help readers to understand characteristics of the nucleosome structural variants that we report in this study.

## Introduction

The nucleosome, composed of eight core histones (H2A_2_, H2B_2_, H3_2_, and H4_2_) and 144~147 bp of DNA, is the fundamental structural unit of chromosomes. X-ray crystal structures of nucleosomes reconstituted *in vitro* with specific nucleosome-positioning sequences revealed that DNA wraps around the core octameric histones in a left-handed direction (Luger et al., 1997; McGinty and Tan, 2015). However, many structural variants of nucleosomes can exist *in vitro* (Lavelle and Prunell, 2007) (Fig. 1A): hexasomes, di-tetrasomes, reversomes (right-handed octameric nucleosome), overlapping dinucleosomes, split nucleosomes, and hemisomes (Arimura et al., 2012; Bancaud et al., 2007; Furuyama et al., 2013; Kato et al., 2017; Zlatanova et al., 2009; Zou et al., 2018). Even within octameric left-handed nucleosomes assembled on the high-affinity nucleosome positioning sequence, Widom 601 (Lowary and Widom, 1998), form multiple structural arrangements of histone octamers, which can be observed by cryo-electron microscopy (cryo-EM) (Bilokapic et al., 2018a, 2018b). Depending on the sequence, the physical length of DNA on a nucleosome can vary (Chua et al., 2012), and DNA can locally stretch at different superhelical locations (SHLs) in nucleosome crystal structures (McGinty and Tan, 2015). Nucleosome binding proteins can also affect the nucleosome structure (Bednar et al., 2017; Dodonova et al., 2020; Falk et al., 2015; Farnung et al., 2017; Liu et al., 2020; Sanulli et al., 2019). While these various nucleosome structures were identified *in vitro* using reconstituted nucleosomes, current reconstructions of *in vivo* nucleosomes are lower than 20 Å resolution (Cai et al., 2018a; Chicano et al., 2019; Eltsov et al., 2018; Scheffer et al., 2011). Cryo-electron tomography (cryo-ET) analyses even suggested that the canonical nucleosome structure may not be abundant in the fission yeast nucleus (Cai et al., 2018b). Thus, the structural heterogeneity of nucleosomes in chromosomes remains unknown (Fig. 1A).

**Figure 1.**
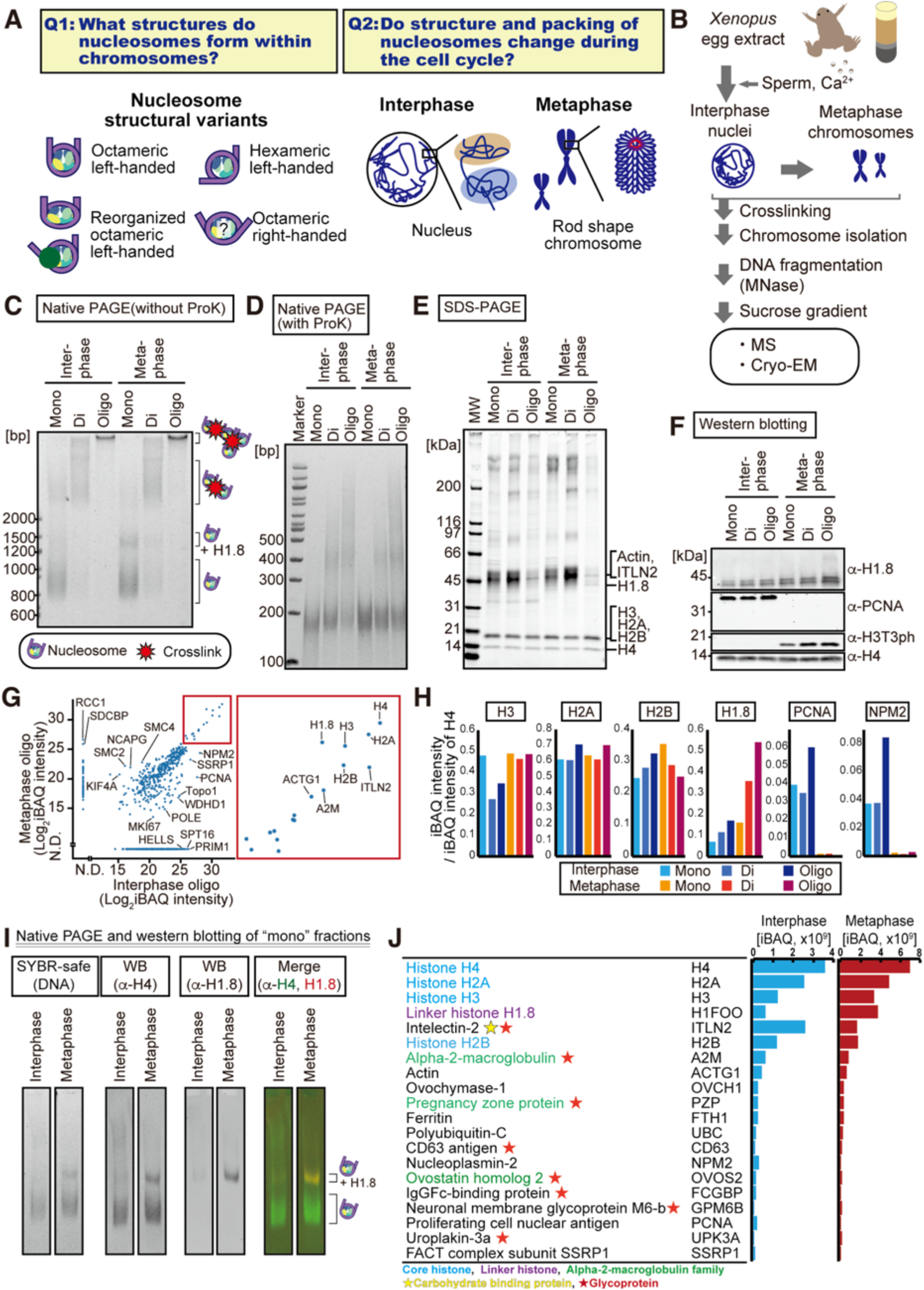
Isolation of nucleosomes from interphase and metaphase chromosomes. **A**, Scopes of this study. **B**, Schematic of nucleosome isolation protocol from *Xenopus* egg extracts. **C, D**, Native PAGE analysis of nucleosome fractions isolated from sperm chromosomes in *Xenopus* egg extracts, without (C) or with (D) proteinase K treatment. Equal DNA amounts were loaded in each fraction. DNA was stained with SYBR-safe. **E,** SDS-PAGE analysis of nucleosomes isolated from sperm chromosomes. Proteins were stained by gel code blue. **F,** Western blot analysis of **E**. **G**, Scatter plot of MS-detected proteins in the oligo nucleosome fractions. Right panel shows a magnified view of the high signal region encased in red. **H**, MS signal intensities of representative proteins, normalized to the H4 signal in each fraction. **I**, Linker histone-bound mono-nucleosomes were detected by native PAGE and western blot. **J,** Protein abundance in the isolated oligo-nucleosome fractions detected by MS. Enrichment of intelectin-2 and glycoproteins (ovochymase and alpha2-macroglobulin family proteins) may reflect abundant O-linked N-acetylglucosamine on chromosomes (Gagnon et al., 2015; Kelly and Hart, 1989).

Chromosome morphology dramatically changes upon entry into mitosis; relatively diffuse interphase chromosomes become individualized and form rod-shaped, compact chromatids (Fig. 1A). Although the DNA loop-forming activity of condensin is the major driver behind the formation of individualized mitotic chromatids (Goloborodko et al., 2016; Hirano and Mitchison, 1994), chromosomes depleted of condensin still exhibit mitosis-specific compaction, suggesting that the nucleosome may contribute to mitotic chromosome compaction (Gibcus et al., 2018; Samejima et al., 2018). Several lines of evidence indicate that nucleosomes are more densely packed in mitosis than in interphase. Nucleosomes assembled *in vitro* on Widom 601 DNA with histones purified from mitosis tend to aggregate more easily than those assembled with histones from interphase (Zhiteneva et al., 2017), and a cryo-ET study suggested that mitotic chromatin is more crowded than interphase chromatin (Cai et al., 2018b). It has been suggested that mitotic deacetylation of the H4 N-terminal tail, which promotes inter-nucleosome interaction through binding to the H2A-H2B acidic patch, promotes local chromatin compaction (Kruitwagen et al., 2015; Vaquero et al., 2006; Wilkins et al., 2014; Zhiteneva et al., 2017). Additionally, electron tomography with DNA labeling (ChromEMT) revealed that the nucleosome concentration in M phase reaches 40-60% (Vo/Vo) (Ou et al., 2017), comparable to the concentration of nucleosomes in a pure crystal (45-55%) (Arimura et al., 2018; Luger et al., 1997). Although a variety of mitosis-specific histone modifications are known (Wang and Higgins, 2013), it remains unclear if the nucleosome structure changes to control mitotic chromatin compaction.

Two major models of nucleosome packing have been proposed: the ordered “30-nm fiber” model and the disordered, “polymer-melt” model (Luger et al., 2012; Maeshima et al., 2010). In the ordered model, nucleosomes on the same linear DNA strand interact together to form a 30-nm fiber, which can be reconstituted *in vitro* (Dorigo et al., 2004; Ekundayo et al., 2017; Robinson et al., 2006; Schalch et al., 2005; Song et al., 2014). This *in vitro* 30 nm-fiber formation is facilitated by the linker histone H1 (Song et al., 2014), which also promotes local chromatin compaction and heterochromatin formation *in vivo* (Fan et al., 2005; Hergeth and Schneider, 2015; Lu et al., 2009). The polymer-melt model was based on evidence that 30-nm fibers cannot be detected by cryo-EM or small angle X-ray scattering in the interphase nuclei and mitotic chromosomes of HeLa cells (Eltsov et al., 2008; Maeshima and Eltsov, 2008; Nishino et al., 2012). In this model, nucleosomes on the same or a different DNA strand can randomly and dynamically interact with each other, creating a highly disordered interdigitated state, although recent genome-wide nucleosome interaction mapping analyses indicate that short nucleosome stretches may locally cluster into ordered structures during interphase (Ohno et al., 2019; Risca et al., 2017). Folding patterns of such ordered nucleosome clusters can vary with the linker DNA angles and lengths of each nucleosome (Grigoryev, 2018). However, direct visualization of linker DNA angles in chromosomes has been technically difficult, making it challenging to speculate on *in vivo* nucleosome packing modes using data obtained by indirect nucleosome-interaction analyses.

Recent technological advances in single particle cryo-EM have enabled structure determination of proteins and protein complexes up to ~2 Å resolution, even of samples lacking the purity required for conventional X-ray crystallography (Cheng, 2018). In principle, structure determination of endogenous proteins should be possible (Kastritis et al., 2017; Nguyen et al., 2015; Yi et al., 2019), but to the best of our knowledge, no direct structural comparisons of protein isolated from different stages of the cell cycle have been reported. Combining the *Xenopus* egg extract system, which enables isolation of large quantities of functional chromosomes at specific cell cycle stages (Funabiki and Murray, 2000; Hirano and Mitchison, 1994), mass spectrometry (MS), and cryo-EM single-molecule analysis, we developed a strategy to determine high-resolution structures of nucleosomes in interphase and metaphase chromosomes. We reveal structural signatures of nucleosomes in chromosomes, as well as their cell cycle-dependent structural variations.

## Results

### Nucleosome isolation from interphase and metaphase chromosomes

We established a method to isolate nucleosomes from functional interphase and metaphase chromosomes while preserving intact protein-DNA structures for cryo-EM analysis (Fig. 1B). *Xenopus* egg extracts arrested at meiotic metaphase II by cytostatic factor (CSF) were released into interphase by adding calcium together with *Xenopus* sperm nuclei, mimicking fertilization (Murray, 1991). Nucleosomes rapidly assembled onto sperm chromosomes, the nuclear envelope formed, and chromosomes were replicated. Adding fresh metaphase-arrested CSF extracts to the interphase extracts induced entry into metaphase, promoting chromosome compaction and spindle formation (Fig. 1B, Fig. S1A) (Shamu and Murray, 1992). Chromosomes in interphase and metaphase extracts were crosslinked with formaldehyde to preserve nucleosome structures, isolated via centrifugation, and then fragmented with micrococcal nuclease (MNase). DNA quantification showed that the majority (60% - 90%) of DNA was solubilized from chromatin with MNase (Fig. S1B). The solubilized nucleosomes were separated by sucrose density gradient centrifugation, yielding fractions of mono-nucleosomes, di-nucleosomes, and larger oligo-nucleosomes (Fig. S1C, S1D, S1E). However, the length of DNA isolated from these complexes was largely equivalent to those from mono-nucleosomes (150-180 bp) (Fig. 1C, 1D, and S1E), indicating that inter-nucleosome DNA linkers were cleaved by MNase. Therefore, the isolated di- and oligo-nucleosome complexes represent clusters of mono-nucleosomes that are chemically crosslinked together.

Biochemical characteristics of the solubilized nucleosome fractions were assessed by western blot and MS. The mitosis-specific histone H3 phosphorylation at Thr3 (H3T3ph) was observed in isolated metaphase nucleosomes but not in interphase nucleosomes (Fig. 1F). Known chromatin proteins, such as proliferating cell nuclear antigen (PCNA) and condensin subunits (CAPD2, SMC2, CAPG, SMC4), showed their expected cell cycle-dependent enrichment on nucleosome fractions (Fig. 1F-H and S1F). Therefore, our nucleosome-isolation method maintained cell cycle-specific biochemical characteristics of chromatin.

These analyses also revealed the preferential association of H1.8 (also known as B4, H1M, and H1Foo), the dominant linker histone variant in *Xenopus* eggs (Dworkin-Rastl et al., 1994; Wühr et al., 2014), with metaphase nucleosomes, rather than interphase nucleosomes (Fig. 1G, 1H, 1J). On native polyacrylamide gel electrophoresis (PAGE) gels, the metaphase mono-nucleosome fraction showed an additional slower-migrating band corresponding to the chromatosome (Fig. 1C, S1D), which contains core histones and H1.8 (Fig. 1I). H1.8 is most enriched in the oligo- and di-nucleosome fractions (Fig. 1H), consistent with the notion that linker histone H1 compacts chromatin *in vitro* and thus leads to increased nucleosome-nucleosome contacts (Renz et al., 1977). While H1.8 accumulation in interphase nuclei have been observed using immunofluorescence (Maresca et al., 2005), our data suggest that chromatosomes containing H1.8 are much more prevalent in metaphase than in interphase.

### Unbiased reconstruction of nucleosome structures in chromosomes by cryo-EM

To examine the diversity of *in vivo* nucleosome structures, we first randomly picked particles from the cryo-EM micrographs containing MNase-solubilized nucleosome fractions and determined all structures that could be reconstructed at an intermediate resolution and in an unbiased manner. Selection bias can propagate due to template-based particle picking and class selection after 2D classification, so we used software tools that did not require any initial templates (Fig. 2A, S2, S3). Specifically, particles were picked from interphase and metaphase oligo-nucleosome fraction micrographs using the CryoSPARC blob picker and were split into two groups based on the resolution and effective classes assigned (ECA) value of the 2D classification (Fig. 2A top, S2) (Punjani et al., 2017). Multiple 3D maps were generated from both groups by CryoSPARC *ab initio* reconstruction (Punjani et al., 2017), resulting in two types of references: 3D maps derived from particles with high-resolution features, and “decoy” references derived from the particles that were noisy, low-resolution, or ambiguous (corresponding to more than two ECA). All picked particles and 3D maps were then used for multireference 3D classification by CryoSPARC (Fig S3A, S3B). This classification method is reminiscent of “random-phase” 3D classification (Gong et al., 2016) and multireference 3D classification with decoy 3D references (Nguyen et al., 2019). However, in our case, instead of using random-phased 3D references or decoy 3D references that are structurally dissimilar cryo-EM maps, we used multiple data-derived decoy 3D references to filter out noisy or low-resolution particles, thereby improving classification accuracy and map quality (Fig. 2A middle).

**Figure 2.**
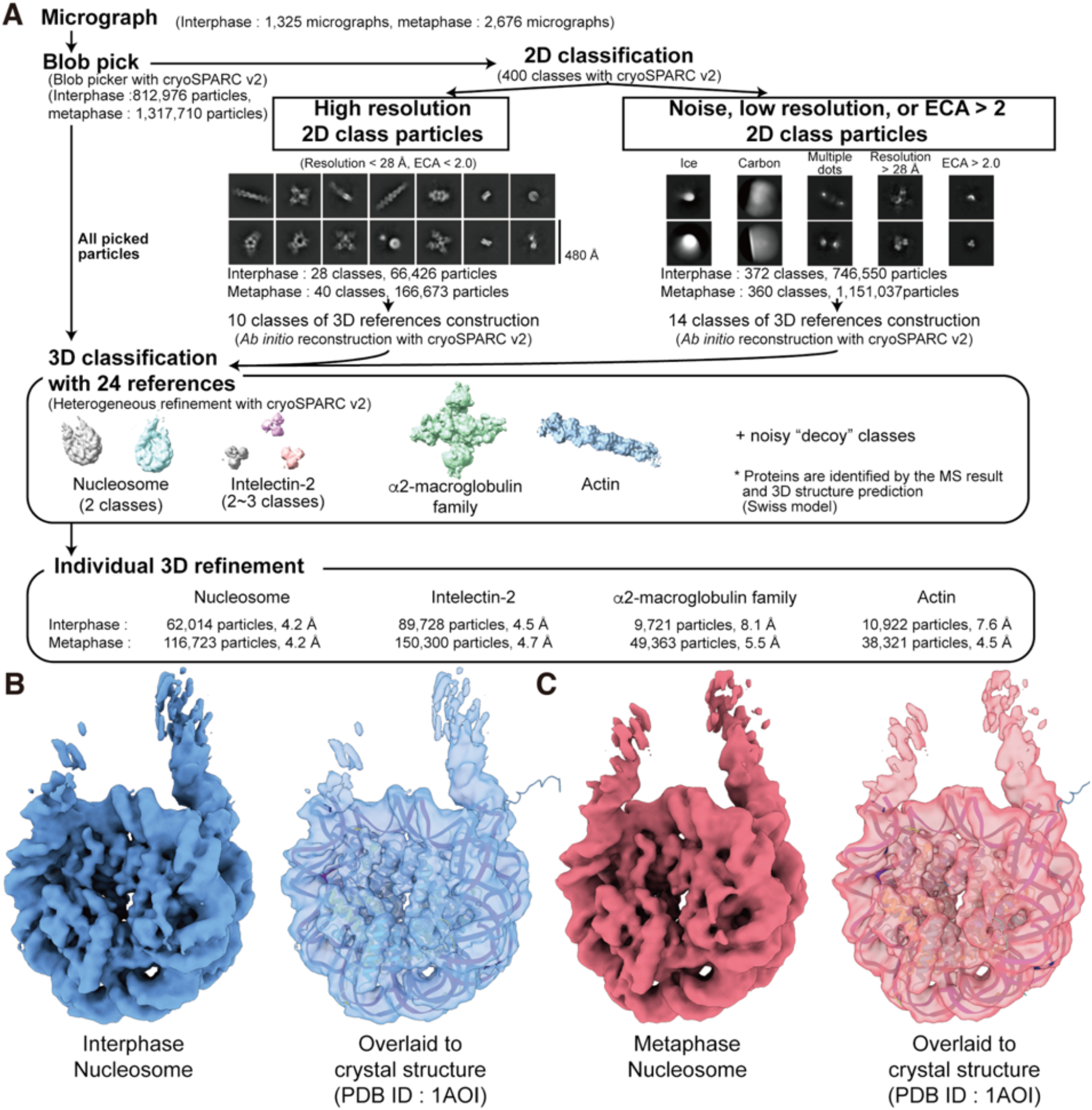
Unbiased cryo-EM analysis revealed the major form of nucleosomes *in vivo.* **A**, Unbiased cryo-EM analysis pipeline. 2D classification panels and 3D references shown were obtained from the metaphase oligo-nucleosome fraction. **B, C**, Reconstructed structures of the interphase nucleosome (B) and metaphase nucleosome (C), and their superposition onto the crystal structure of the canonical left-handed octameric nucleosome (PDB ID: 1AOI).

Using this decoy classification method, four distinct protein complex structures were obtained simultaneously (Fig. 2A middle, S3A, S3B). Guided by the MS-based protein abundance (Fig. 1J) and sequence-based 3D structure modeling (Waterhouse et al., 2018), the structures were identified as an octameric mono-nucleosome, intelectin-2 (25~70% abundance relative to histone H4), alpha2-macroglobulin family protein (10~20 % of H4), and actin (5~7% of H4). Both interphase and metaphase nucleosome structures aligned well with the crystal structure of the canonical left-handed octameric nucleosome (Fig. 2B, 2C, S3C, S3D) (Luger et al., 1997). While cryo-EM maps of actin and intelectin matched published EM and crystal structures, respectively (Fujii et al., 2010; Wangkanont et al., 2016), major structural changes were seen in our alpha2-macroglobulin cryo-EM structure compared to the recombinant human alpha2-macroglobulin crystal structure (Fig. S3E-G) (Marrero et al., 2012), likely because the intracellular alpha2-macroglobulin in egg extracts should represent its inactive form. Thus, our pipeline successfully reconstructed multiple structures at once without the need for any template-picking or 2D class selection-based approaches, which were previously required to determine multiple structures from crude samples (Kastritis et al., 2017; Kyrilis et al., 2019; Verbeke et al., 2018, 2020).

Our unbiased reconstructions indicate that on average, nucleosomes in both interphase and metaphase chromosomes exist in the canonical left-handed octameric structure (Fig. 2B and C). While it is conceivable that the CryoSPARC algorithms failed to reconstruct unstable nucleosome structural variants, it is unlikely that these variants were selectively broken during the grid-freezing step, since free nucleosome-size DNA fragments were rarely observed on the micrograph under our optimized freezing conditions (Fig. S2B).

### High-resolution cryo-EM structures of interphase and metaphase nucleosomes

The nominal resolution of chromosomal nucleosome structures obtained by the unbiased reconstruction method was 4.2 Å (Fig. 2). The limitation in this method was caused by the random particle picking process using the CryoSPARC blob picker with a 100 Å minimum inter-particle distance, which excludes particles in closely-spaced nucleosome clusters, reducing the number of picked particles. To further improve resolution by picking more nucleosome-like particles in a comprehensive manner, we employed Topaz, a convolutional neural network-based particle picking program with positive-unlabeled learning (Bepler et al., 2019), which we first trained on manually picked particles (Fig. S4A, particle picking). Octameric nucleosome-like 2D classes were then manually selected to generate initial 3D references (Fig. S4A, initial 3D reference reconstruction). Using an octameric nucleosome-like 3D map, along with four decoy maps that were generated from non-nucleosome-like particles, 3D decoy classification of all Topaz picked particles was performed with CryoSPARC (Fig. S4A, 3D decoy classification). After 3D refinement with Relion-3.1, structures of interphase and metaphase nucleosomes in the oligo-nucleosome fractions were reconstructed at a nominal 3.4 and 3.5 Å resolution, respectively (Fig. 3A, 3B, and S4B). These high-resolution maps allowed us to build atomic models of the interphase and metaphase nucleosomes with clearly resolved amino acid side chains (Table S2). For example, the aromatic side chain of phenylalanine 98 of H2A.X-F1/F2, the predominant H2A isoform in egg (Shechter et al., 2009), could be clearly visualized and distinguished from the leucine or isoleucine residues of other H2A isoforms (Fig. S5A). The root-mean-square distance (R.M.S.D.) measurements demonstrated that the histone structures in atomic models for interphase and metaphase nucleosomes are essentially identical to those of X-ray crystal structures of nucleosomes assembled *in vitro* with specific nucleosome positioning sequences (Fig. 3C, 3D), with one notable exception.

**Figure 3.**
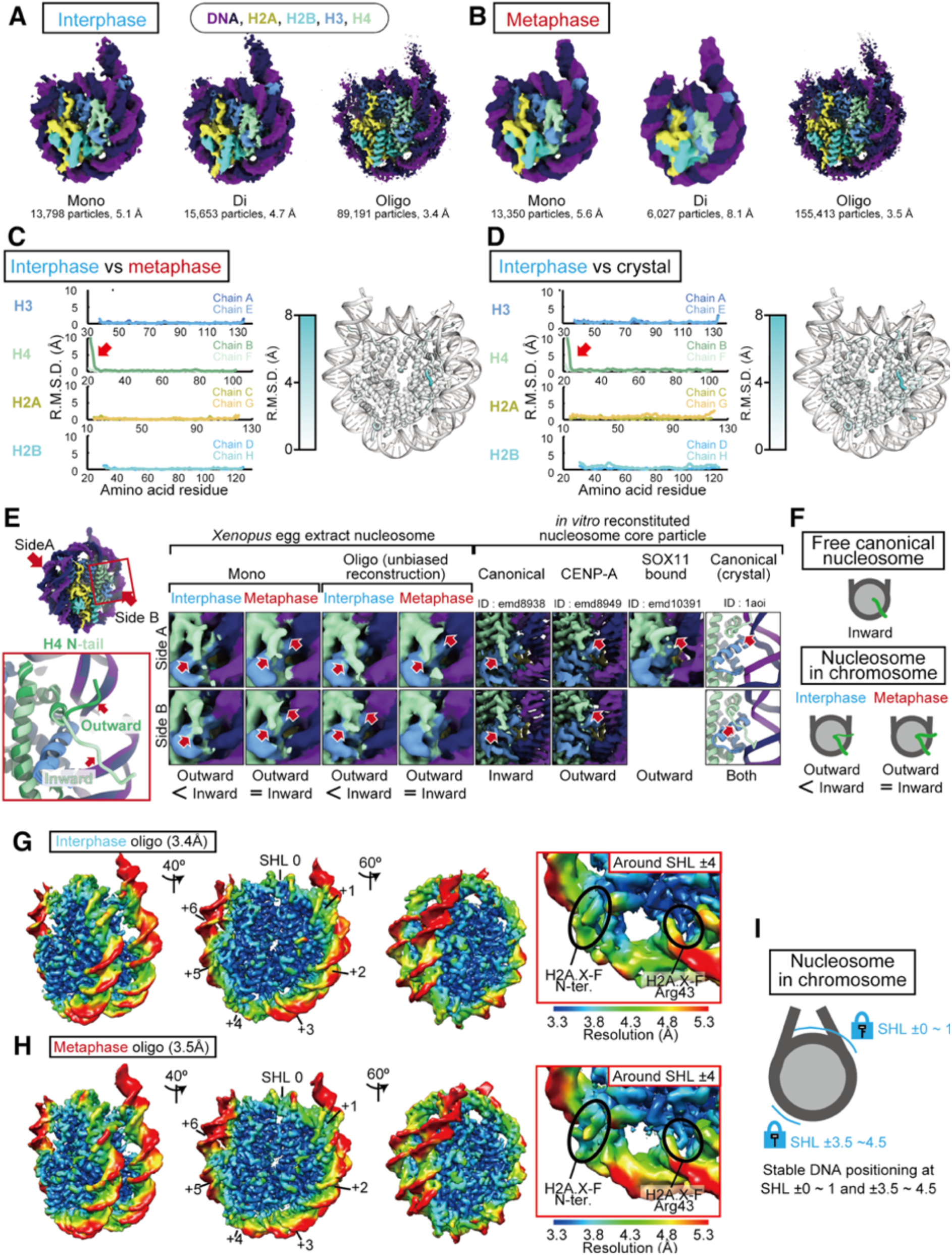
High-resolution cryo-EM structures of interphase and metaphase nucleosomes. **A**, **B,** Cryo-EM structures of the nucleosome core particle from interphase (A) and metaphase (B) chromosomes. Maps were sharpened with the Postprocess function of Relion 3.1. **C, D.** Structural comparisons between interphase and metaphase nucleosomes (C) or between interphase nucleosomes and the canonical crystal structure of the nucleosome (D) (PDB ID: 1AOI). Each histone protein chain in the atomic coordinates are named separately following the definition in the nucleosome crystal structure (PDB ID: 1AOI) (Luger et al., 1997). Root-mean-square deviation (R.M.S.D.) for all Cα atoms for each of the structures was calculated using PyMol and plotted in the specified combinations (left), or R.M.S.D. values for all Cα atoms were mapped onto the atomic coordinates of the interphase oligo fraction nucleosome, and the 3D structure were drawn and colored using PyMol (right). **E,** Comparisons of the H4 N-terminal region. Left, schematic defining “Side A”, “Side B”, and “inward” and “outward” H4 N-terminal orientations. Right, Cryo-EM maps of the H4 N-terminal region, segmented and colored using atomic coordinates determined in this study. **F,** Graphical explanation of H4 N-terminal tail structure in chromosomal nucleosome. **G, H**, Local resolution maps of interphase (G) and metaphase (H) nucleosomes. Left, local resolution maps of the entire nucleosome. Right, local resolution maps of DNA around SHL ±3.5~4.5. Local resolution was calculated with BLOCRES and BLOCFILT in the BSOFT package. **I.** Graphical explanation of DNA positioning in chromosomal nucleosome.

A unique structural difference between interphase and metaphase nucleosomes can be seen at the base segment of the H4 N-terminal tail (Fig. 3C, arrow). Several different structures of *in vitro* reconstituted nucleosomes have revealed two distinct “outward” and “inward” orientations of this region (Fig. 3E, left). Cryo-EM structures of the canonical mono-nucleosome show an inward H4 N-terminal tail orientation (Zhou et al., 2019), whereas the nucleosome with histone H3 variant CENP-A and the nucleosome bound to the transcription factor SOX11 exhibit an outward orientation (Ali-Ahmad et al., 2019; Arimura et al., 2019; Dodonova et al., 2020; Zhou et al., 2019). These structural differences suggest that the H4 N-terminal tail of *in vitro* reconstituted mono-nucleosomes is mainly oriented inward, although the orientation can be changed to outward through incorporation of histone variants or binding of additional proteins (Fig. 3E). In the canonical nucleosome crystal, both inward and outward orientations can exist (Fig. 3E right), suggesting that the outward orientation also can be formed in the mono-nucleosome within a highly concentrated and packed environment. In our reconstructions, both interphase and metaphase nucleosomes had a mixed density of inward and outward conformations. However, in interphase, nucleosomes preferentially formed the inward conformation, while in metaphase, both inward and outward conformations were observed at a comparable level (Fig. 3E left). Altogether, this observation suggests that the H4 N-terminal tail can change its orientation between the inward and outward conformations within chromosomes, and mitosis-specific changes in nucleosome-nucleosome or nucleosome-binding partner interactions may shift the H4 N-terminal tail orientation (Fig. 3F).

### DNA at specific superhelical locations is exceptionally ordered in chromosomal nucleosomes

Despite heterogeneity of DNA sequences, our high-resolution metaphase and interphase nucleosome maps exhibit extraordinary homogeneity in DNA structure, such that the DNA backbone is clearly delineated (Fig. 3A, 3B). At the center of the nucleosomal DNA, a single base pair aligns at the nucleosome dyad (Flaus et al., 1996), whose location in the nucleosome structure is defined as SHL 0 (Klug et al., 1980). In crystal structures, other DNA segments on the nucleosome can stretch, depending on the DNA sequence and other bound proteins (Chua et al., 2012; McGinty and Tan, 2015). Strikingly, in our high-resolution cryo-EM structures of nucleosomes isolated from interphase and metaphase chromosomes, DNA segments at SHL ±3.5~4.5 as well as SHL ±0~1 exhibited lower B-factor values (Fig. S5B) and higher local resolutions (Fig. 3G and 3H) compared to other regions. Despite non-uniform DNA sequence and length, the EM density resolution of DNA around these segments was high enough to distinguish positioning of phosphates (Fig. 3G and 3H). This indicates that, in chromosomes, DNA is more stably positioned at SHL ±0~1 and SHL ±3.5~4.5 than at other DNA segments. Consistent with this observation, the number of DNA-histone interactions formed with basic amino acid residues on histones is particularly high in these regions (Fig. S5C-E). At SHL ±3.5~4.5, amino acids 15 to 44 of H2A.X-F interact with the negatively-charged DNA backbone via basic residues (Arg18, Lys21, Arg30, Arg33, and Arg43) (Fig. 5SE, S5F), which are highly conserved among species and H2A variants (Fig. S5G). These data indicate that the histone moieties and DNA around SHL ±0~1 and SHL ±3.5~4.5 of *in vivo* nucleosomes are exceptionally stable, independent of DNA sequence and cell cycle stages, while other regions, such as SHL ±2~3, ±5~6 and linker DNA, are relatively flexible (Fig. 3I).

### Unbiased identification of nucleosome structural variants

We next examined the heterogeneity of nucleosomes in chromosomes with the recently developed the 3D variability analysis (3DVA) in CryoSPARC (Punjani and Fleet, 2020), which models cryo-EM data into multiple series of 3D structures (motion frames), each representing a potential “trajectory” of motion (Fig. 4A, Movie S1). Importantly, each trajectory represents a principal component of the variance present in each dataset, so each 3DVA component presented here are unbiased representations of the major structural variation. Three prominent apparent structural variations were observed (Fig. 4A, black, blue, and orange arrows, Movie S1).

**Figure 4.**
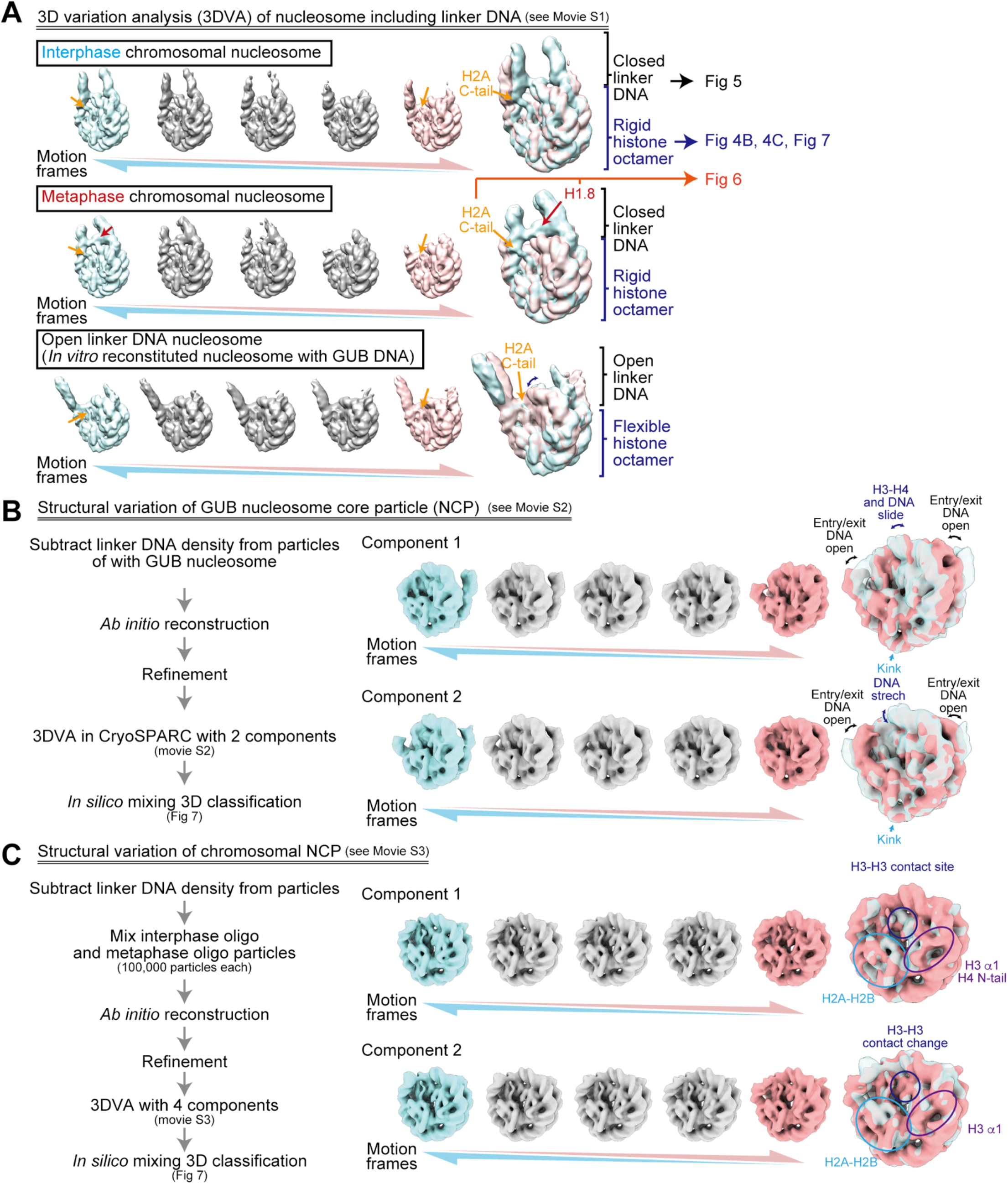
3D variability analysis of the nucleosomes. **A**, 3D variability analysis of the nucleosomes containing linker DNA. Five representative structures (motion frame 1, 5, 9, 13, 17) from 3DVA component 1 class (total 20 frames) are shown. The structures of motion frames 1 and 17 were selected and overlayed for each nucleosome. EM densities for the H2A C-terminal region (orange arrows) and H1.8 (red arrows) are shown. See Movie S1 for full dataset. **B, C**, NCP structural variation of GUB nucleosome (B) and chromosomal nucleosome (C). Five representative structures (motion frames 3, 7, 11, 15, 19) from 3DVA component 1 and 2 classes (total 21 frames each) are shown. Structures of motion frame 3 and 19 were selected and overlayed for each nucleosome.

First, variations in the linker DNA angle were detected (Fig. 4A black arrow and brackets, Movie S1). A previous study using small-angle scattering analysis suggested that linker DNA angles may differ by DNA positioning sequence (Yang et al., 2011). Indeed, linker DNA angles of the nucleosome assembled on the nucleosome positioning sequence, GUB (Adams and Workman, 1995; An et al., 1998), were much more open than those of nucleosomes assembled on 601 DNA (Fig. 4A lower panel and S7F). To our surprise, despite the heterogeneity of DNA sequences, linker DNA angles of interphase and metaphase nucleosomes were generally closed, relative to *in vitro*-reconstituted nucleosomes with GUB DNA (Fig. 4A). Quantitative assessment of this observation will be described in Figure 5.

**Figure 5.**
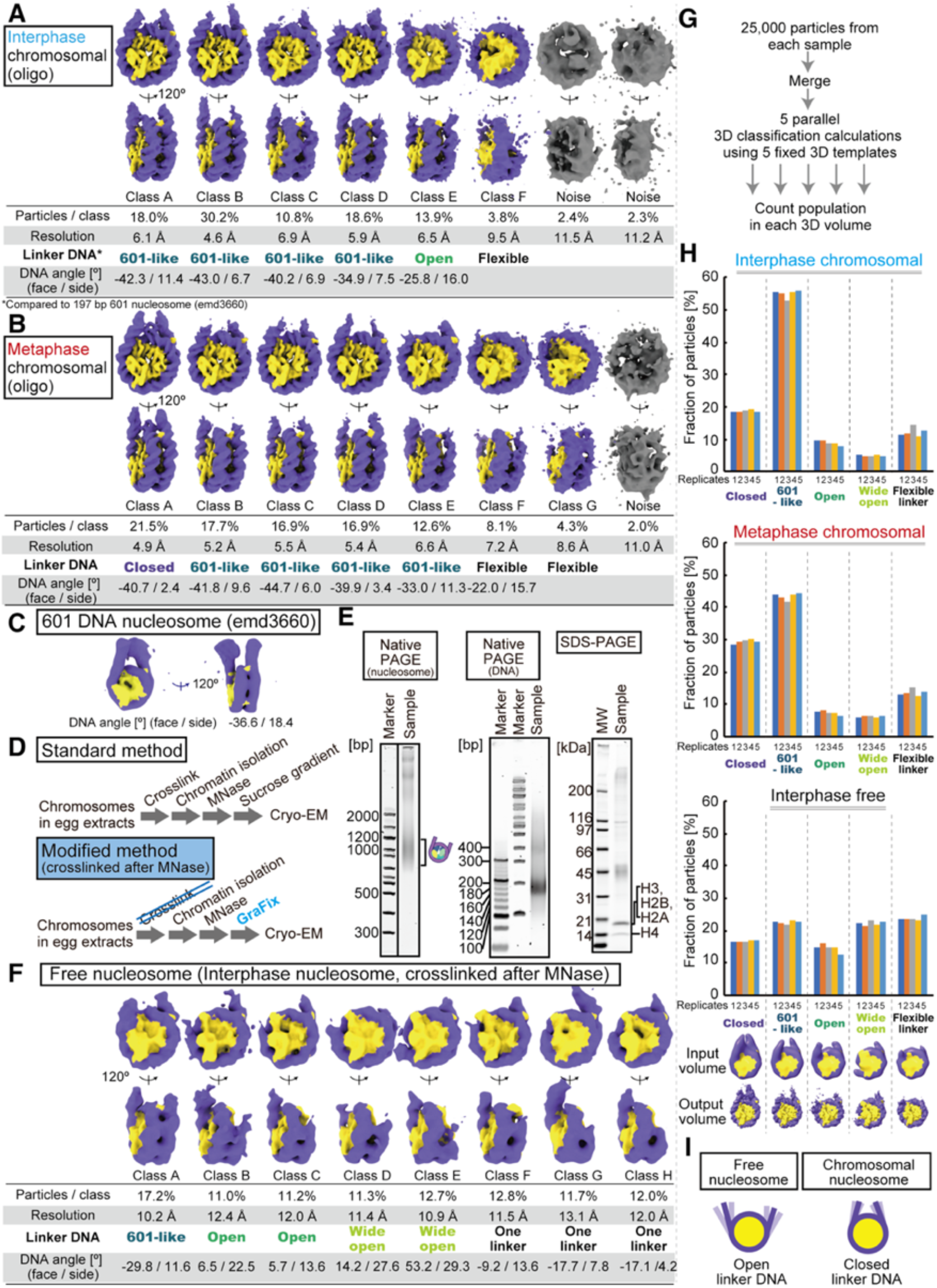
The linker DNA angles of a majority of interphase and metaphase nucleosomes are closed. **A, B,** 3D structure classes of nucleosomes in the oligo-nucleosome fractions of interphase chromosomes (A), and metaphase chromosomes (B). To classify the linker DNA angle of each class, the cryo-EM structure of the 601 DNA nucleosome (ID : emd3660) (Bednar et al., 2017) was used as a standard for comparisons (*Closed*: Linker DNA does not fit into the linker DNA of the 601 nucleosome and the extra density is observed inward of the 601 linker DNA density; *601-like*: Linker DNA density fits into the linker DNA of 601 nucleosome; *Open*: Linker DNA does not fit into the linker DNA of the 601 nucleosome, extra density is observed outside of the 601 linker density, and the H3 N-terminal region (a.a. 37-56) still binds DNA at the nucleosome entry/exit site; *Wide open*: Linker DNA does not fit into the linker DNA of the 601 nucleosome, extra density is observed outside of the linker DNA density of the 601 nucleosome, and the H3 N-terminal region (a.a. 37-56) does not bind the nucleosome entry/exit site DNA.; *Flexible* : No visible linker DNA density on either end of the nucleosomal DNA.; *One linker*: No visible linker DNA density on one end of the nucleosomal DNA). **C,** Cryo-EM structure of the nucleosome with 197bp 601 DNA (ID : emd3660) (Bednar et al., 2017), which was used as a reference for the linker DNA angle. **D,** Schematic of the modified method to isolate nucleosomes crosslinked after chromatin fragmentation. **E,** Native PAGE and SDS-PAGE of the purified nucleosomes crosslinked after chromatin fragmentation. **F,** 3D structure classes of nucleosomes crosslinked after chromatin fragmentation. **G**, Schematic of the *in silico* mixing 3D classification pipeline. **H**, *In silico* mixing 3D classification with fixed 3D references for the oligo fractions of interphase nucleosomes, the oligo fractions of metaphase nucleosomes, and the nucleosomes crosslinked after chromatin fragmentation. 3D classifications were performed with five different linker DNA angle 3D maps low-pass filtered to 32 Å. The relative fractions assigned to each class of the five independent 3D classification analyses are shown. 3D map images at the bottom indicate the input and output of the 3D classifications. **I,** Graphical summary of Figure 5.

Second, extra cryo-EM density was observed in chromosomal nucleosomes, particularly when linker DNA angle was most closed (Fig. 4A, Movie S1, orange and red arrows). For both interphase and metaphase, the densities of the H2A C-terminal tail reached DNA at the DNA entry/exit site of the nucleosome as the DNA closed inward (Fig. 4A, Movie S1, orange arrows). In addition, as the linker DNA angle closed, an additional cryo-EM density appeared at the nucleosome dyad position, particularly in metaphase nucleosomes (Fig. 4A, Movie S1, red arrows). High resolution analysis of these extra cryo-EM densities will be described in Figure 6.

**Figure 6.**
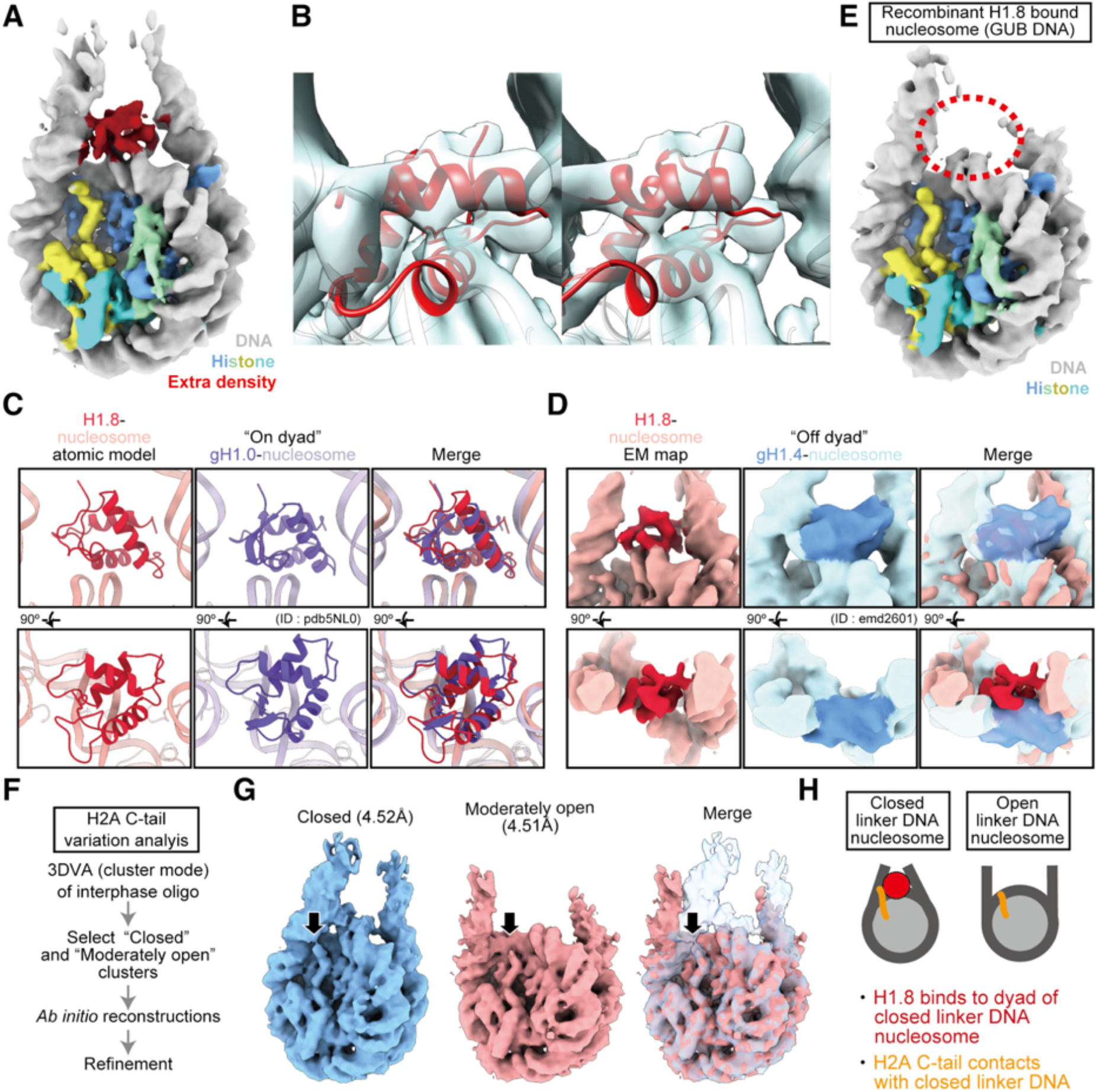
Linker histone H1.8 binds on the nucleosome dyad in metaphase chromosomes. **A**, Cryo-EM structure of the nucleosome class with an extra density (in red) on the dyad isolated from the oligo fraction of metaphase nucleosomes. **B**, Stereo images of the cryo-EM map superimposed on an atomic model of a H1.8-bound nucleosome (red). **C**, The atomic model of the H1.8-bound nucleosome superimposed on the “on-dyad” H1-bound nucleosome crystal structure (PDB ID: 5NL0). **D**, The cryo-EM map of the H1.8-bound nucleosome superimposed on the “off-dyad” H1 bound nucleosome cryo-EM map (emd ID: 2601). **E,** Cryo-EM structure of the *in vitro* reconstituted H1.8-bound nucleosome with GUB DNA. **F**, Schematic of the reconstruction of “closed” and “relatively open” linker DNA nucleosome maps. **G**, H2A C-tail density in the “closed” and “moderately open” linker DNA nucleosome maps. Black arrows indicate H2A C-tail. **H**, Graphical explanation of the linker DNA angle dependent observation of H1.8 and H2A C-tail-DNA interaction.

Third, major structural rearrangements of nucleosome core particles (NCPs) are seen, particularly in GUB nucleosomes (Fig. 4A blue arrows and brackets, Movie S1). To focus on the NCPs structural variation, we employed the 3DVA analysis on GUB NCP and chromosomal NCP after subtracting linker DNA signal from the original particles (Fig. 4B, 4C, Movie S2). This analysis revealed two different types of NCP structural rearrangement. The first type was the major NCP rearrangement, which was characterized by diverse NCP surface outlines seen in 3DVA of GUB NCPs (Fig. 4A-B, Movie S1-3, also see Fig. 7A). The second type is the relatively minor NCP rearrangement observed only after subtraction of linker DNAs in 3DVA of chromosomal NCPs (Fig. 4C, Movie S2-3, also see Fig. 7B). In these minor structural variants, rearranged orientations of histone α-helices and the H4 N-terminal tail can be seen (Movie S3). Population analysis of these structural variants will be described in Figure 7.

**Figure 7.**
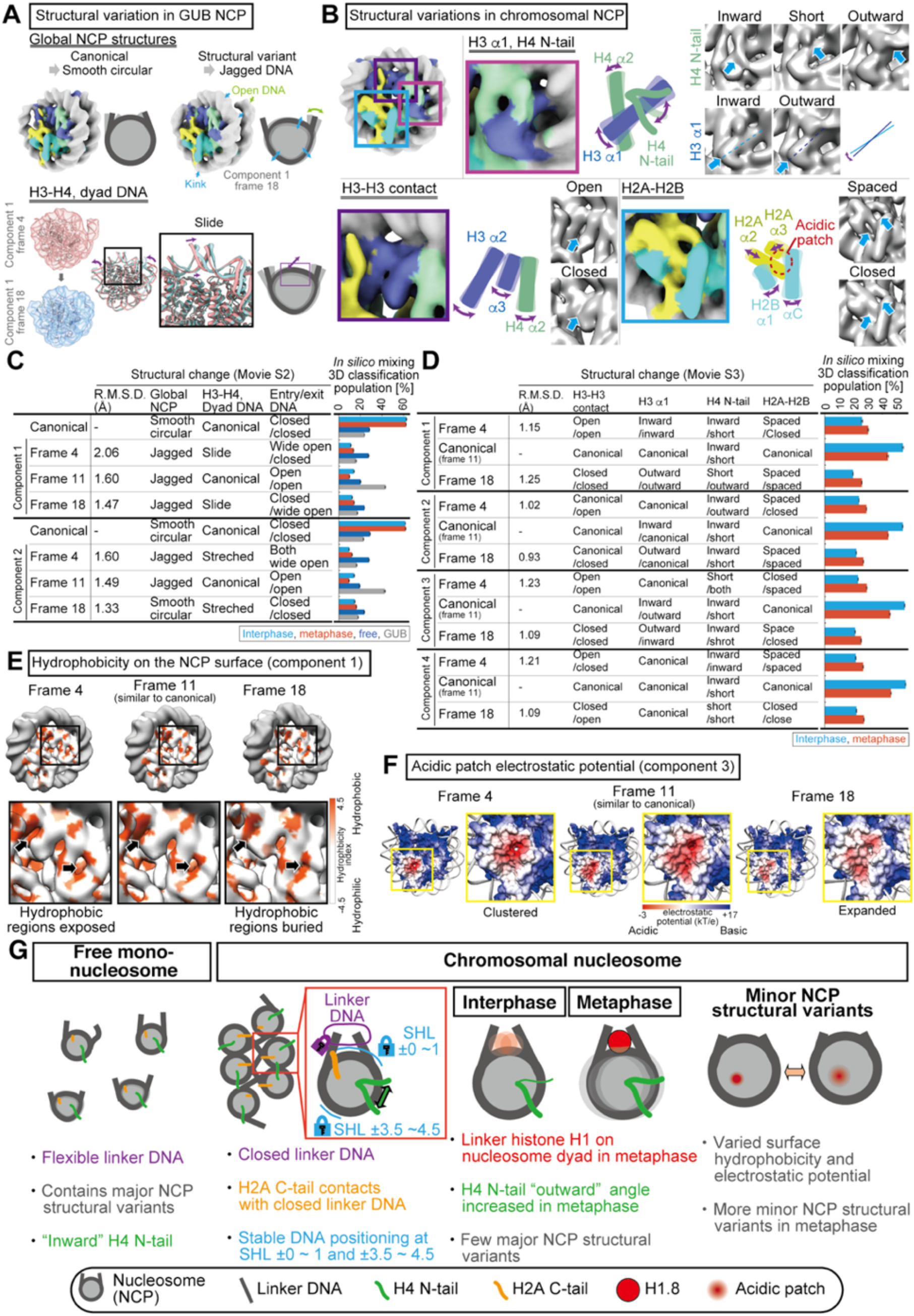
Structural variation of chromosomal NCP and their cell cycle-dependent changes. **A,** Types of major NCP structural variations in GUB NCP 3DVA. **B,** Types of minor NCP structural variations in chromosomal NCP 3DVA. **C**, *In silico* mixing 3D classification of major NCP variants. Means and standard deviations of the relative fractions assigned to each class of the five parallel *in silico* mixing 3D classification analyses are shown. **D**, *In silico* mixing 3D classification of minor NCP variants. Means and standard deviations of the relative fractions assigned to each class of the five parallel *in silico* mixing 3D classification analyses are shown. **E**, Surface hydrophobicity of the chromosomal NCP structural variants. Hydrophobicity indexes were mapped on the EM maps of the representative time frame of 3DVA component 1 of chromosomal NCP. **F**, Electrostatic potential of the chromosomal NCP structural variants. Surface electrostatic potentials for the atomic models for the representative time frame of 3DVA component 3 of chromosomal NCPs were shown. **G**, Structural characteristics of free mono-nucleosomes and nucleosomes isolated from interphase or metaphase chromosomes.

### Linker DNAs are closed in the majority of resolved nucleosome structures in interphase and metaphase chromosomes

To quantitatively assess the structural variations of linker DNA angles observed using 3DVA, we used *ab initio* reconstruction to generate 6-8 nucleosome 3D references representing distinct linker DNA structural classes, and then classified all nucleosome-like particles (Fig. 5A, B, S6A, B, Table S3). We compared the location and angle of linker DNAs of each of these structural classes with those of cryo-EM nucleosome structures assembled on the strong nucleosome positioning sequence Widom 601 (Fig. 5C, S6C) (Bednar et al., 2017). Linker DNA angles in the majority of interphase and metaphase nucleosomes were equivalent to, or more closed inward than, those of nucleosomes assembled on Widom 601 DNA (Fig. 5A, B and S6A, B, E).

To test if the abundance of nucleosomes with closed linker DNA angles reflects structural characteristics of nucleosomes in chromosomes, rather than biased reconstruction of nucleosomes with closed linker DNA angles, we also determined cryo-EM structures of chromosomal nucleosomes using a modified protocol in which native nucleosomes were first released from interphase chromosome by MNase and then crosslinked (Fig. 5D, 5E, 5F). The majority of these “free” mono-nucleosome structures were classified into structures with open linker DNA angles that were wider than those assembled on the Widom 601 sequence (Fig. 5F). These results suggest that there is an active mechanism to close the linker DNA of nucleosomes in chromosomes, but once mono-nucleosomes are released from chromosomes by MNase prior to fixation, linker DNAs can adopt more flexible open conformations, as seen in the GUB nucleosomes.

Since *ab initio* structure reconstruction might have failed to generate relatively minor structural classes and resulted in skewed class distributions, we next performed *in silico* mixing 3D classification (Hite and MacKinnon, 2017): we combined particles from the three consensus refinements (interphase nucleosomes, metaphase nucleosomes, and interphase nucleosomes crosslinked after MNase treatment), and conducted a new round of classification analysis using five manually-selected 3D references with distinct linker DNA angles (Fig. 5G, H). Five parallel classification calculations yielded consistent results: around 70% of interphase or metaphase nucleosomes were assigned to classes with a similar or more closed linker DNA angle relative to the 601 nucleosome (Fig. 5H top and middle), while these proportions were reduced in free mono-nucleosomes fixed after MNase digestion (Fig. 5G bottom). 3D reconstructions of each class confirmed that the linker DNA angle differences were preserved through the classification process (Fig. 5H bottom, EM maps). Although the precise proportions of each class vary based on classification methods (Fig. 5A, B, F, H), both analyses support the conclusion that the majority of nucleosomes in chromosomes possess closed linker DNAs (Fig. 5I), and nucleosomes with closed linker angles are more prevalent in metaphase chromosomes than in interphase chromosomes (Fig. 5H, S6D, S6E).

### Linker histone H1.8 binds on the nucleosome dyad in metaphase chromatin

As noted above, an extra dyad density reminiscent of the linker histone was observed in the metaphase nucleosome with closed linker DNA (Fig. 4A, 5B, Movie S1). Further 3D classification of Class A shown in Fig. 5B enabled structure determination at 4.4 Å resolution (Fig. 6A, S7A-C). Since histone H1.8 preferentially cofractionated with metaphase nucleosomes (Fig. 1G-J), we built an initial atomic model of the H1.8-bound nucleosome through homology modeling using a published chromatosome crystal structure containing the globular domain of histone H1.0 (gH1.0) (Bednar et al., 2017). The initial model was docked onto the cryo-EM map and refined. The extra EM density reasonably corresponded to our refined H1.8 atomic model (Fig. 6B). Previously, structural analyses of *in vitro* reconstituted chromatosomes and molecular dynamics simulations suggested that linker histones may bind to the nucleosome in two distinct modes; “on-dyad”, where H1 binds at the center of the nucleosome DNA (Bednar et al., 2017; Zhou et al., 2015), and “off-dyad”, where H1 binds more than 5 bp off-center of the nucleosome DNA (Adhireksan et al., 2020; Song et al., 2014; Woods and Wereszczynski, 2020) Since on-dyad H1 interacts with both linker DNAs and constricts them, while off-dyad H1 provides linker DNAs with more flexibility, it has been proposed that on-dyad H1 more effectively compacts chromatin than off-dyad H1 (Zhou et al., 2015). The H1.8 position in our cryo-EM structure of the metaphase chromatosome more closely corresponds to the on-dyad structure than the off-dyad structure (Fig. 6C, 6D, 6H) (Bednar et al., 2017; Zhou et al., 2015). In addition, the Class A structure where H1.8 density could be visualized also had stronger density for the linker DNAs (Fig. 5B), consistent with the idea that linker histone on-dyad stabilizes the linker DNA (Bednar et al., 2017).

Initially, we did not expect to resolve H1.8 on mitotic nucleosomes, since the EM density of H1.8 was barely seen when reconstituted *in vitro* using purified recombinant proteins with 176 bp GUB DNA (Fig. 6E, S7E-H) (Adams and Workman, 1995; An et al., 1998). Unlike nucleosomes assembled on Widom 601 sequence, linker DNA angles were open in most of the GUB nucleosomes (Fig. S7F). The recombinant H1.8 stoichiometrically bound to the GUB nucleosome (Fig. S7G), and closed the linker DNA of the GUB nucleosome, even though no clear H1.8 EM density was observed (Fig. S7H). This suggests that an active mechanism exists on mitotic chromosomes to stabilize H1.8 at the on-dyad position.

In addition to the extra EM density corresponding to H1.8, 3DVA analysis revealed density for the H2A C-terminal tail in nucleosomes with closed linker DNAs (Fig. 4A, Movie S1). To better assess this structural correlation, we generated 3D maps of interphase nucleosomes with closed and open linker DNAs at comparable resolutions (4.51 Å and 4.52 Å) (Fig. 6F, 6G). The cryo-EM density corresponding to the H2A C-terminal tail is stronger and continuous with the DNA density in the nucleosome with closed linker DNAs, while it is weaker in the nucleosome with open linker DNAs (Fig. 6G, black arrows), suggesting a putative role for the H2A C-terminal tail in influencing the linker DNA angle (Fig. 6H).

### Structural variants of the NCP in chromosome and their cell cycle-dependent change

Our 3DVA analysis revealed two different types of NCP structural variants: the major NCP structural variants that are linked to large linker DNA angle variations (Fig. 7A, and also see Fig. 4A, 4B, Movie S1-2) and the minor NCP structural variants that are not correlated with linker DNA angle variations (Fig. 7B, and also see Fig 4C, Movie S3). The major structural variants are characterized by jagged NCP outlines with sliding of the H3-H4 tetramer, as compared to the smoother circular shape of the canonical NCP (Fig. 7A). To test if the major NCP variants can be seen in chromosomal nucleosomes, we employed *in silico* mixing 3D classification for each 3DVA component using four reference 3D maps; one representing the canonical nucleosome structure (generated from composite particles of interphase and metaphase), and three representing the major nucleosome structural variants (picked from 3DVA motion frames of GUB NCP, which showed major NCP rearrangement; see Methods) (Fig. 7A, C). While more than 70% of NCPs were assigned to the major NCP structural variants when nucleosomes had been freed from chromatin by MNase before fixation (“Free” in Fig. 7C), only ~40 % of the interphase and metaphase NCPs were assigned to the major NCP structural variants when chromatin had been fixed prior to MNase (Fig. 7C). Consistent with this result, structural variations of the NCP were observed in the free nucleosome even without removing the linker DNA density (Movie S1). These results suggest that formation of the major NCP structural variants related to linker DNA angle variations is suppressed in chromosomes, relative to free mono-nucleosomes.

The minor NCP structural variants are characterized by rotation of H3 α1 and reorientation of the H4 N-terminal tail (Fig. 7B, magenta), opening and closing of two H3 α-helices at the dyad (violet), and opening and closing of the H2A-H2B acidic patch (light blue). *In silico* mixing 3D classification for the minor NCP rearrangement revealed that these minor NCP structural variants are more frequently observed in metaphase than in interphase (Fig. 7D). As seen in averaged chromosomal nucleosome structures (Fig. 3E), variants with the outward H4 N-terminal tail were more prevalent in metaphase than in interphase (3DVA component 1, frame 18; component 2, frame 4; and component 3, frame 4). In some of these variants, buried hydrophobic amino acid residues were exposed to the NCP surface (Fig. 7E), and there is redistribution of the electrostatic potential at the H2A-H2B acidic patch (Fig 7E, 7F), suggesting that these structural rearrangements of NCPs may affect proteins binding on NCPs or nucleosome-nucleosome interactions. Altogether, these results suggest that nucleosomes in chromosomes are more structurally homogeneous than free mono-nucleosomes, but structural changes in the NCP can still occur, with greater frequency in mitotic chromosomes than in interphase chromosomes (Fig. 7G).

## Discussion

### Characteristics of nucleosome structures in interphase and metaphase chromosomes

Here we present near-atomic resolution structures of nucleosomes from interphase nuclei and metaphase chromosomes. On average, reconstructed nucleosome-like structures isolated from both interphase and metaphase chromosomes are almost identical to previously reported cryo-EM and crystal structures of *in vitro* reconstituted canonical left-handed octameric nucleosomes assembled on defined nucleosome positioning sequences (Fig. 2, 3) (Chua et al., 2016; Luger et al., 1997). Although it is still possible that minor populations of nucleosomes form alternative structures, we propose that our reconstructions represent the major forms in mitotic and interphase chromosomes for the following reasons. First, to avoid the possibility that unstable nucleosome structural variants were selectively broken during purification, we optimized a gentle purification procedure and cryoprotectant buffer (Fig. S2B), enabling successful structure determination of the five most abundant protein complexes (nucleosome, chromatosome, alpha2-macroguloblin, intelectin-2, actin filament) detected by MS (Fig. 1J, 2). While this procedure was able to simultaneously classify and determine these highly divergent structures from crude samples, the only nucleosome-like structure reconstituted using an unbiased *ab initio* structure-determination pipeline was the octameric left-handed nucleosome (Fig. 2, S3A, S3B). Second, around 60-90 % of chromosomal DNA was solubilized after MNase digestion (Fig. S1B), suggesting that the solubilized nucleosomes analyzed by cryo-EM did not represent minor fractions of chromatin. Third, structures of *in vitro* reconstituted mono-nucleosomes, and nucleosomes crosslinked after chromatin fragmentation, were more heterogenous than those of chromosomal nucleosomes (Fig. 5I, Movie S1), suggesting that our procedures enabled preservation and detection of dynamic histone and DNA movements. Altogether, our data suggest that the major nucleosome structure is the octameric left-handed nucleosome, and that global histone architectures are generally homogeneous within chromatin, regardless of DNA sequence or cell cycle stage.

3D structure variability analysis showed that closed linker DNA angle is highly correlated with visibility of the H2A C-terminal tail and H1.8 (Fig. 4A, 6A, 6G, Movie S1). H1.8 density was only visible on nucleosomes with the most-closed linker DNA angle, in agreement with the previous observation that H1 binding constricts the linker DNA angle (Bednar et al., 2017). In addition, our data suggest that the H2A C-terminal tail may regulate linker DNA conformation (Fig. 7F). Since the C-terminal tail of H2A is poorly conserved between paralogs and is a target of various modifications (Corujo and Buschbeck, 2018), it is tempting to speculate that linker DNA conformation is regulated by these H2A variants and modifications. Consistent with this idea, recent cryo-EM study showed that the linker DNA is open in the nucleosome reconstituted with the H2A variant H2A.Z.2.2, which has a flexible C-terminal tail (Zhou et al., 2020).

Although the polymer-melt model proposed that nucleosomes are not packed in an ordered manner (Eltsov et al., 2008; Maeshima and Eltsov, 2008; Nishino et al., 2012), the polymer-melt status of chromatin may not solely consist of random distribution of mono-nucleosomes, but may also contain “nucleosome motifs,” comprised of specific oligo-nucleosome arrangements (Krietenstein and Rando, 2020; Ohno et al., 2019; Ricci et al., 2015; Risca et al., 2017). Nucleosomes with closed linker DNA angles in our cryo-EM structures may be a signature of such “nucleosome motifs.” However, it is possible that the prevalence of closed linker DNA angles is specific to *Xenopus* egg and early embryos, where zygotic *de novo* transcription is suppressed due to its specialized chromatin status (Amodeo et al., 2015; Newport and Kirschner, 1982a, 1982b).

Local EM resolution analysis suggests that DNA segments at SHL −1 to +1 and SHL ± 4 are structurally more stabilized than other DNA segments (Fig 3G-I). Since the H2A.X-F amino residues 15-44 that interact with DNA at SHL ±4 are highly conserved among H2A variants across different species, stable positioning of those two DNA segments is likely to be a common, intrinsic feature of nucleosomes (Fig. S5G). Stabilization at these two specific DNA segments may explain why RNA polymerase II halts around SHL ±5 and SHL ±1 on the nucleosome during transcription *in vivo* and *in vitro* (Kujirai et al., 2018; Weber et al., 2014). In contrast, the local resolution at SHL ± 2~3.5 and ± 5~6 was relatively low, suggesting that DNA at these sites may slide on the nucleosome more easily (Fig. 3G, 3H). The ATPase module of many chromatin remodeling factors, such as BAF, RSC, ISWIa, Chd1, INO80, and SWR1, binds to SHL ±2.5 or ±6, implying that chromatin remodeling factors are optimized to target the flexible DNA regions on the nucleosome (Ayala et al., 2018; Eustermann et al., 2018; Farnung et al., 2017; Han et al., 2020; He et al., 2020; Liu et al., 2017; Willhoft et al., 2018; Yan et al., 2019; Ye et al., 2019). It is also possible that the DNA stability at SHL ±1 and SHL ± 4 is necessary for the directional sliding of DNA.

NCP structural variants have been reported by cryo-EM and biochemical assays using *in vitro* reconstituted nucleosomes (Bilokapic et al., 2018a, 2018b; Falk et al., 2015; Sanulli et al., 2019), but their existence and biological significance *in vivo* were unknown. Using 3DVA and *in silico* mixing 3D classification, we revealed two distinct types of NCP rearrangements in reconstituted recombinant nucleosomes and in chromosomal nucleosomes: the major structural variants coupled to diverse linker DNA angles, and the minor structural variants that are seen independently of linker DNA angle (Fig. 4B, 4C, 7B, Movie S2, S3). Our data suggest that major NCP variants, prevalent in free mono-nucleosomes, are suppressed in chromosomes (Fig. 4, 7, Movie S1-3), while minor NCP structural variants commonly exist in chromosomes (Fig. 4C, 7B, Movie S3). Since structures of mono-nucleosomes that were fixed after being freed from chromosomes by MNase digestion have more major structural variants (Fig. 7C), local chromatin environment may limit the structural flexibility of nucleosomes.

### Implications for models of chromatin compaction during the cell cycle

Hirano hypothesized that condensin drives chromatin compaction and suppresses DNA unwinding by converting (−) crossings of linker DNAs to (+) crossings, which is inhibited in interphase by linker histones and released in M phase by the phosphorylation of linker histones (Hirano, 2014). However, our cryo-EM analysis did not reveal mitotic conversion of linker DNAs. The majority of interphase and metaphase nucleosomes contained closed linker DNAs (Fig. 5, S6 and Movie S1), while histone H1.8 preferentially associates with the metaphase nucleosome at the on-dyad position and stabilizes (−) crossing (Fig. 6A). The preferential association of H1.8 to mitotic nucleosomes may contribute to local chromatin compaction, since the linker histone C-terminal disordered region may act as a liquid-like glue for chromatin, and linker histone-containing poly-nucleosomes form condensed and less mobile chromatin (Gibbs and Kriwacki, 2018; Gibson et al., 2019; Turner et al., 2018). It is also possible that H1 on the nucleosome dyad may regulate condensin activity, as condensin prefers to bind nucleosome-free DNA (Kong et al., 2020; Zierhut et al., 2014).

Since most histone chaperones and nucleosome remodelers tend to dissociate from mitotic chromatin (Funabiki et al., 2017; Jenness et al., 2018), it was surprising that minor NCP structural variants revealed by 3DVA and *in silico* mixing 3D classification were more prevalent in metaphase than in interphase nucleosomes (Fig. 7B). One of the most prominent variations was the orientation of the H4 N-terminal tail; the outward orientation was more prevalent in metaphase nucleosomes than in interphase nucleosomes (Fig. 3E, Fig. 7B, D). In the X-ray crystallography of recombinant mono-nucleosomes, inward H4 N-terminal tails mediate the inter-nucleosomal binding via interactions with the acidic patch (Luger et al., 1997). From cryo-EM structures of recombinant oligo-nucleosomes, it can be speculated that outward H4 N-terminal tails also can mediate the inter-nucleosomal binding (Song et al., 2014). In chromosomes, this outward orientation may be correlated with the enhanced interaction between the H4 tail and the acidic patch of the adjacent nucleosome (Kruitwagen et al., 2015). In addition, many interspaces on the NCP surface were newly generated in the minor NCP structural variants (Fig 7E, Movie S3), mimicking a situation where nucleosome binding of CENP-C and Swi6 increased solvent accessibility of buried histone residues (Falk et al., 2015; Sanulli et al., 2019). Exposure of these buried histone residues changes the hydrophobicity and electrostatic potential on the nucleosome surface (Fig. 7E, 7F), potentially affecting specificity and affinity of nucleosome-binding targets (Skrajna et al., 2020). Exposure of hydrophobic surfaces may enhance weak, transient multivalent nucleosome-nucleosome interactions (Sanulli et al., 2019), while a change in charge distribution at the acidic patch may affect binding targets, including the H4 N-terminal tail. Thus, the population-level changes of these minor structural variants during the cell cycle may reflect global changes in nucleosome-interacting proteins and the chromatin compaction mechanism. Future studies are needed to test the role of these cell cycle-dependent regulations of structural variations of the nucleosome.

In this work, we were able to successfully determine the structures of nucleosomes in functional chromosomes at near-atomic resolution, and quantitatively analyze their structural variations. Since our method is compatible with chromatin immunoprecipitation (ChIP) (Wal and Pugh, 2012), it may be adapted for determination of nucleosome structures with specific chemical modifications and/or nucleosome-binding factors. Future improvement of this strategy may capture high-resolution structures of protein-DNA complexes while they are functioning on chromosomes *in vivo*.

## Acknowledgments

We are grateful to Seth Darst, James Chen, Elizabeth Campbell, Gregory Alushin, Hiroshi Suzuki, Mark Ebrahim, Johanna Sotiris, and Honkit Ng for their technical advice and assistance for the Cryo-EM. We thank Hiroshi Kimura and Pavan Choppakatla for sharing reagents, and Soeren Heissel for MS analysis. We thank Haruka Oda and Coral Zhou for their technical advice for chromatin isolation from *Xenopus* egg extract. We thank Seth Darst, James Chen, Gregory Alushin, Viviana Risca, Hitoshi Kurumizaka, Tomoya Kujirai, Risa Fujita, Kayo Nozawa, Yoshimasa Takizawa, Christian Zierhut, Simona Giunta, and Ryan Kim for their critical comments to the manuscript. This work was conducted with the help of Rockefeller High Performance Computing resource center, Rockefeller Proteomics Resource Center, The Evelyn Gruss Lipper Cryo-Electron Microscopy Resource Center, Rockefeller Electron Microscopy Resource Center. This work was funded by National Institutes of Health Grants R35GM132111 to H.F. Y.A. was supported by JSPS Overseas Research Fellowships.

## Author Contributions

Y.A. conducted experiments using *Xenopus* egg extract. R.M.S. conducted the experiment using *in vitro* reconstituted nucleosome under the guidance of Y.A. Y.A. and R.M.S. performed cryo-EM analysis. R.F. developed the decoy-based 3D classification. Y.A. and H.F. designed the experiments and wrote the paper. R.M.S. and R.F. edited the paper. H.F. oversaw the entire project.

## Legends

**Movie S1. The 3D variation analysis of the nucleosomes. Related to Figure 4, 5, 6, 7.**

3D motion movie of the nucleosomes calculated with 3DVA in CryoSPARC. For all samples, the number of 3DVA components was set to two and both components are shown. Movies were created with Chimera using all output 3D map frames of 3DVA. EM densities for H2A C-terminal region (orange arrows) and H1.8 (red arrows) are shown.

**Movie S2. The 3D variation analysis of GUB NCP. Related to Figure 4, 7.**

3D motion movie of the GUB NCP calculated with 3DVA in CryoSPARC. The number of 3DVA components was set to two and both components are shown. Major NCP structural variation sites are pointed out with arrows.

**Movie S3. The 3D variation analysis of chromosomal NCP. Related to Figure 4, 7.**

3D motion movie of the chromosomal NCP calculated with 3DVA in CryoSPARC. The number of 3DVA components was set to four and all components are shown. Minor NCP structural variation sites are shown in zoomed-in panels.

## Data and Code Availability

Cryo-EM maps will be available after publication of the peer-reviewed paper as EMD-22797 (nucleosome in interphase mono nucleosome fraction), EMD-22798 (nucleosome in interphase di nucleosome fraction), EMD-22790 (nucleosome in interphase oligo nucleosome fraction), EMD-22800 (nucleosome in metaphase mono nucleosome fraction), EMD-22801 (nucleosome in metaphase di nucleosome fraction), EMD-22791 (nucleosome in metaphase oligo nucleosome fraction), EMD-22792 (H1.8 bound nucleosome in metaphase oligo nucleosome fraction), EMD-22799 (nucleosome in interphase nucleosome crosslinked after MNase), EMD-22903 (reconstituted nucleosome with GUB DNA), EMD-22802 (H1.8 bound reconstituted nucleosome with GUB DNA), EMD-22793 (nucleosome structure of interphase oligo nucleosome fraction with unbiased reconstruction), EMD-22796 (nucleosome structure of metaphase oligo nucleosome fraction with unbiased reconstruction), EMD-22794 (alpha2-macroglobulin structure of interphase oligo nucleosome fraction with unbiased reconstruction), and EMD-22796 (alpha2-macroglobulin structure of metaphase oligo nucleosome fraction with unbiased reconstruction). Atomic models are available in the PDB for 7KBD (nucleosome in interphase oligo nucleosome fraction), 7KBE (nucleosome in metaphase oligo nucleosome fraction), and 7KBF (H1.8 bound nucleosome in metaphase oligo nucleosome fraction).

## Materials and Methods

### *Xenopus* egg extracts and chromatin formation

Cytostatic factor (CSF) metaphase-arrested *X. laevis* egg extracts were prepared with the method described previously (Murray, 1991). To prevent spontaneous cycling of egg extracts, 0.1 mg/ml cycloheximide was added to the CSF extract. For interphase chromosome preparation, *X. laevis* sperm nuclei (final concentration 2000/μl) were added to 8 ml of CSF extracts, which were incubated for 90 min at 20 °C after adding 0.3 mM CaCl_2_, which releases CSF extracts into interphase. To monitor spindle assembly, Alexa594-labeled-bovine brain tubulin (final concentration 19 nM) was added to the extract during the incubation. For metaphase sperm chromosome preparation, Cyclin B Δ90 (final concentration 24 μg/ml) and 4 ml fresh CSF extract was added to 8 ml of extract containing the interphase sperm nuclei prepared with the method described above. The extracts were incubated 60 min at 20 °C, during which each tube was gently mixed every 10 min. Animal husbandry and protocol approved by Rockefeller University’s Institutional Animal Care and Use Committee were followed.

### Nucleosome isolation from *Xenopus* egg extracts chromosomes

To crosslink the *Xenopus* egg extracts chromosomes, nine times volume of ice cold buffer XL (80 mM PIPES-KOH [pH 6.8], 15 mM NaCl, 60 mM KCl, 30 % glycerol, 1 mM EGTA, 1 mM MgCl_2_, 10 mM β-glycerophosphate, 10 mM sodium butyrate, 2.67 % formaldehyde) was added to the interphase or metaphase extract with chromosomes, which were further incubated for 60 min on ice. These fixed samples were layered on 3 ml of buffer SC (80 mM HEPES-KOH [pH 7.4], 15 mM NaCl, 60 mM KCl, 1.17 M sucrose, 50 mM glycine, 0.15 mM spermidine, 0.5 mM spermine, 1.25x cOmplete EDTA-free Protease Inhibitor Cocktail (Roche), 10 mM β- glycerophosphate, 10 mM sodium butyrate, 1 mM EGTA, 1 mM MgCl_2_) in 14 ml centrifuge tubes (Falcon, #352059) and spun at 3,300 (2,647 rcf) rpm at 4 °C for 40 min using JS 5.3 rotor (Beckman Coulter) in Beckman Avanti J-26S. Chromatin was collected from the bottom of the centrifuge tube. Collected chromatin was further spun through buffer SC as described above. Chromatin was collected from the bottom of the centrifuge tube, and then this step was repeated once again. Only at the 1^st^ round of centrifugation over sucrose cushion, chromatin trapped at the boundary between the extract and buffer SC was also collected and applied for 2^nd^ round centrifugation over sucrose cushion. The isolated interphase nuclei and metaphase chromosomes were associated with white aggregations and spindle microtubules, respectively. To remove them, chromosomes were gently pipetted with a 2 ml pipette. These chromosomes were pelleted by centrifugation at 5,492 rpm (2,500 rcf) using SX241.5 rotor (Beckman Coulter) in Allegron X-30R. The chromosome pellets were resuspended with 200 μL of buffer SC. To digest chromatin, 1.5 U/μL of MNase (Worthington Biochemical Corporation) and CaCl_2_ were added to 7.4 mM, and the mixture was incubated at 4 °C for 6 h. MNase reaction was stopped by adding 900 μL MNase stop buffer (15 mM HEPES-KOH [pH 7.4], 150 mM KCl, 5 mM EGTA, 10 mM β-glycerophosphate, 10 mM sodium butyrate, 5 mM DTT). The soluble fractions released by MNase were isolated by taking supernatants after centrifugation at 13,894 rpm (16,000 rcf) at 4 °C for 30 min using SX241.5 rotor in Allegron X-30R (Beckman Coulter). The supernatants were collected and layered onto the 10-22 % linear sucrose gradient solution with buffer SG (15 mM HEPES-KOH [pH 7.4], 50 mM KCl, 10-22 % sucrose, 10 μg/ml leupeptin, 10 μg/ml pepstatin, 10 μg/ml chymostatin, 10 mM sodium butyrate, 10 mM β-glycerophosphate, 1 mM EGTA) and spun at 32000 rpm (124,436 rcf) and 4 °C for 13 h using SW55Ti rotor (Beckman Coulter) in Beckman Optima L80. The samples were fractionated from the top of the sucrose gradient.

To prepare the interphase nucleosome that was crosslinked after MNase, *Xenopus* egg extracts chromosomes was diluted with the buffer XL without formaldehyde and was incubated for 60 min on ice. After the chromatin isolation and MNase treatment with the same method as described above, chromatin was isolated and fixed using the gradient fixation (GraFix) method (Kastner et al., 2008). Briefly, a sucrose gradient was created using 2.3 ml of buffer GF1 (15 mM HEPES-KOH [pH 7.4], 50 mM KCl, 22 % sucrose, 10 μg/ml leupeptin, 10 μg/ml pepstatin, 10 μg/ml chymostatin, 10 mM sodium butyrate, 10 mM β-glycerophosphate, 1 mM EGTA, and 4 % paraformaldehyde aqueous solution [EM Grade, Electron Microscopy Sciences)) and 2 ml of buffer GF2 (15 mM HEPES-KOH [pH 7.4], 50 mM KCl, 10 % sucrose, 10 μg/ml leupeptin, 10 μg/ml pepstatin, 10 μg/ml chymostatin, 10 mM sodium butyrate, and 10 mM β-glycerophosphate, 1 mM EGTA). The supernatant of MNase treated chromatin were overlaid onto the sucrose gradient and spun at 32000 rpm (124,436 rcf) for 13 h at 4 °C using SW55Ti rotor (Beckman Coulter) in Beckman Optima L80.

### Native and SDS-PAGE analysis of nucleosomes

For the native PAGE for nucleosome (Fig 1C, 5E), nucleosome fractions containing 100 ng DNA were to load onto a 0.5x TBE 6 % native PAGE gel. For the native PAGE for nucleosomal DNA (Fig 1C, 5E), nucleosome fractions containing 100 ng DNA were mixed with 1 μL of 10 mg/ml RNaseA (Thermo Scientific) and incubated at 55 °C for 30 min. To deproteinize and reverse-crosslink DNA, RNaseA treated samples were then mixed with 1 μL of 19 mg/ml Proteinase K solution (Roche) and incubated at 55 °C for overnight. Samples were loaded to 0.5x TBE 6 % native PAGE. (Figure 1D, 5E). Native PAGE gels were stained by SYBR-safe or SYTO-60 to detect DNA. For SDS-PAGE analysis (Figure 1E), nucleosome fractions containing 200 ng DNA were loaded to a 4-20 % gradient gel.

### Western blotting of native PAGE to detect H1.8-bound nucleosome

Fifteen μL of the interphase-mono and metaphase-mono fractions just after sucrose gradient was applied onto 6 % native PAGE with x0.5 TBE, and DNA was stained by SYBR-safe (invitrogen). After acquiring image, the native PAGE gel was soaked in transfer buffer (25 mM Tris, 192 mM glycine and 20% methanol) and blotted onto the nitrocellulose membrane (GE) using TE42 Tank Blotting Units (Hoefor) at 15 V, 4 °C for 4 h. Histone H4 was detected by 2.4 μg/ml of mouse monoclonal antibody (CMA400) provided by H. Kimura (Hayashi-Takanaka et al., 2015). H1.8 was detected by 1μg/ml of rabbit polyclonal antibody against full-length *Xenopus laevis* H1.8 (Jenness et al., 2018). As secondary antibodies, IRDye 680LT goat anti-rabbit IgG (Li-Cor 926-68021; 1:10,000) and IR Dye 800CW goat anti-mouse IgG (Li-Cor 926-32210; 1:15,000) were used. The images were taken with Odyssey Infrared Imaging System (Li-Cor).

### Nucleosome dialysis and concertation for Cryo-EM and MS

Nucleosome containing sucrose gradient fractions were collected and dialyzed against 600 ml dialysis buffer (10 mM HEPES-KOH [pH 7.4] 30 mM KCl, 1 mM EGTA, 0.3 μg/ml leupeptin, 0.3 μg/ml pepstatin, 0.3 μg/ml chymostatin, 1 mM sodium butyrate, 1 mM β-glycerophosphate) using Tube-o-dialyzer 15 kDa (G-Biosciences) at 4 °C. The samples were further dialyzed twice against the fresh dialysis buffer. The samples were concentrated using Amicon Ultra centrifugal filters 10K (Millipore Sigma). Absorbance at wavelength 260 nm were 2.705, 1.342, 3.756, 2.959, 2.564, 2.06, and 5.244 for interphase mono nucleosome, interphase di nucleosome, interphase oligo nucleosome, interphase nucleosome crosslink after MNase, metaphase mono nucleosome, metaphase di nucleosome, and metaphase oligo nucleosome, respectively. The isolated nucleosomes were stored at 4 °C.

### Mass spectrometry

The samples containing 800 ng DNA were incubated at 95 °C for 30 min to reverse crosslinking. DNA amounts were estimated by the 260 nm absorbance. Samples were applied to SDS-PAGE (4-20% gradient gel) (Bio-Rad). The Gel was stained by Coomassie Brilliant Blue G-250 (Thermo Fisher). Gel bands were excised by scalpel and cut into pieces of approximately 2mm*2mm. Destaining, in-gel digestion and extraction was performed as described (Shevchenko et al., 2007). Extracted peptides were purified using in-house constructed micropurification C18 tips. Purified peptides were analyzed by LC-MS/MS using a Dionex3000 HPLC equipped with a NCS3500RS nano- and microflow pump coupled to a Q-Exactive HF mass spectrometer (Thermo Scientific). Peptides were separated by reversed phase chromatography (solvent A: 0.1% formic acid in water, solvent B: 80% acetonitrile, 0.1% formic acid in water) across a 70-min gradient. Spectra were recorded in positive ion data-dependent acquisition mode, fragmenting the 20 most abundant ions within each duty cycle. MS1 spectra were recorded with a resolution of 60 k and AGC target of 3e6. MS2 spectra were recorded with a resolution of 30 k and ACC target of 2e5. Spectra were queried against a *Xenopus laevis* database (Wühr et al., 2014) (3413 sequences) concatenated with common contaminants using MASCOT through Proteome Discoverer v.1.4 (Thermo Scientific). Oxidation of M and acetylation of protein N-termini were set as variable modifications and carbamidomethylation of C was set as static modification. A false discovery rate of 1% was applied. A large number of identified peptides map to multiple highly similar proteins. iBAQ values of these peaks were summed before drawing the figures (Schwanhüusser et al., 2011).

### Western blotting

Nucleosome fractions containing 200 ng DNA were applied for SDS-PAGE with 4-20 % gradient SDS-PAGE gel (Bio-rad). Proteins were transferred to the nitrocellulose membrane (GE) from the SDS-PAGE gel using TE42 Tank Blotting Units (Hoefer) at 15 V, 4 °C for 4 h. As primally antibodies, 1 μg/ml of Mouse monoclonal H3T3ph antibody 16B2 (Kelly et al., 2010) and 1 μg/ml of mouse monoclonal PCNA antibody (Santa Cruz SC-56) were used. For H4 and H1.8 detection, antibodies described above were used as primally antibodies. As secondary antibodies, IRDye 680LT goat anti-rabbit IgG (Li-Cor 926-68021; 1:10,000) and IR Dye 800CW goat anti-mouse IgG (Li-Cor 926-32210; 1:15,000) were used. The images were taken with Odyssey Infrared Imaging System (Li-Cor).

### Histone preparation for recombinant nucleosome

All histones were purified with the method described previously (Zierhut et al., 2014). Briefly, bacterial expressed *X. laevis* H2A, H2B, H3.2, and H4 were purified from inclusion bodies. Histidine-tagged histones (H2A, H3.2, and H4) or untagged H2B expressed in bacteria were resolubilized from the inclusion bodies by incubation with the 6 M guanidine HCl. For Histidine-tagged histones, the resolubilized histidine-tagged histones were purified with Ni affinity chromatography using Ni-NTA beads (Qiagen). For untagged H2B, resolubilized histones were used for H2A-H2B dimer formation without further purification. To reconstitute H3–H4 tetramer and H2A–H2B dimer, denatured histones were mixed at a concentration of ~45 μM and dialyzed to refold histones with removing the guanidine. Histidine-tags were removed by TEV protease treatment, and H3–H4 tetramer and H2A–H2B dimer were isolated with gel-filtration chromatography on a HiLoad 16/60 Superdex 75 pg column (GE Healthcare). Fractions containing (H3–H4 tetramer and H2A– H2B dimer) were concentrated and stored at −80 °C.

### DNA preparation for recombinant nucleosome

Four tandem 176 bp GUB DNA sequences (An et al., 1998) were cloned into pUC18 vector (pUC18-GUB_176×4). The *E.coli* DH5a that possessing the plasmid was cultured in 1.5x TBG-M9-YE medium with 25 μg/ml carbenicillin at 37 °C for overnight. The plasmid was purified using plasmid plus giga kits (Qiagen) following standard protocol. The 176 bp GUB DNA was cut out from the purified plasmid with EcoRV (New England Biolabs). The plasmid backbone was removed by polyethylene glycol precipitation using PEG-6000. The 176 bp GUB DNA was recovered from the supernatant of the polyethylene glycol precipitation fraction and further purified with gel-filtration chromatography on a HiLoad 16/60 Superdex 75 pg column (GE Healthcare) using TE buffer (10 mM Tris-HCl [pH 7.5] and 0.1 mM EDTA). DNA containing fractions were collected and stored at −20 °C.

The DNA sequence of the 176 bp GUB DNA is the following. 5’-ATCCC TCTAG ACGGA GGACA GTCCT CCGGT TACCT TCGAA CCACG TGGCC GTCTA GATGC TGACT CATTG TCGAC ACGCG TAGAT CTGCT AGCAT CGATC CATGG ACTAG TCTCG AGTTT AAAGA TATCC AGCTG CCCGG GAGGC CTTCG CGAAA TATTG GTACC CCATG GAAGA T-3’

### Recombinant H1.8 purification

N-terminal His-tag, GST, and TEV protease recognition site fused *X. laevis* linker histone H1.8 was expressed in *E. coli* strain C43(DE3) using a pColdII vector (Takara Bio) (pColdII-His-GST-TEVsite-H1.8). *E. coli* C43(DE3) was transformed with pColdII-His-GST-TEVsite-H1.8 and plated onto a carbenicillin containing agar plate. Colonies were incubated overnight at 37 °C and used to inoculate 4 L of TBG-M9 without glucose containing 50 μl/ml of carbenicillin. Bacteria containing flasks were incubated at 37 ° C, 150 rpm for 4 h. Flasks were chilled on ice for approximately 15 min then incubated at 15 °C, 150 rpm, for overnight. Bacteria were spun down at 6000 rpm at 4 °֯C for 30 min in JLA8.1 rotor (Beckman Coulter) in Beckman Avanti J-26S. The bacteria pellet was resuspended in sonication buffer (50 mM Tris-HCl [pH 7.5], 500 mM NaCl, 10% glycerol, 2 mM β-mercaptoethanol, 1 mM PMSF, 5 mM EDTA, 1x cOmplete protease inhibitor cocktail (EDTA-free) (Roche)). Cells were lysed via sonication and pelleted at 30,000 rpm, 4 ° for 45 min in 70Ti rotor in Beckman optima L80. Supernatant was incubated with Glutathione Sepharose 4B resin (GE Healthcare) for 1 h and column washed overnight with GST wash buffer (50 mM Tris-HCl [pH 7.5], 1 M NaCl, 5% glycerol, 2 mM β-mercaptoethanol, 1 mM PMSF, 5 mM EDTA, 0.05x cOmplete protease inhibitor cocktail (EDTA-free). Protein-bound beads were then washed with two column volume of Ni wash buffer (50 mM Tris-HCl [pH 7.5], 100 mM NaCl, 5% glycerol, 10 mM imidazole, 2 mM β-mercaptoethanol). Beads were then incubated with TEV protease at 16 °C for 2 overnights and the Tag removed H1.8 was eluted with Ni wash buffer. Purified H1.8 was concentrated with Amicon Ultra 10K (Millipore) and further purified with HiLoad Superdex 75 16/60 (GE Healthcare) in equilibrium with superdex buffer (10 mM HEPES-KOH [pH 7.4], 30 mM KCl). Fractions containing H1.8 were collected and applied to HiTrap Heparin HP (GE Healthcare) column. The column was washed with superdex buffer and H1.8 was eluted with the linear gradient of KCl (30 mM to 1 M). Fractions containing H1.8 were collected and dialyzed to superdex buffer with Tube-O-DIALYZER (15 kDa) (G-Biosciences). Dialyzed H1.8 was concentrated with Amicon Ultra 10K (Millipore), flash frozen with liquid nitrogen, and stored at −80 °C.

### Recombinant nucleosome reconstitution

Approximately 1.6 μM of purified 176 bp GUB DNA was mixed with 3.2 μM H3-H4 reconstituted histone dimers and 3.5 μM H2A-H2B reconstituted dimers with the salt dialysis method described previously (Zierhut 2014). The sample mixture was transferred into a dialysis cassette and placed into a high salt buffer (10 mM Tris-HCl [pH 7.5], 1 mM EDTA, 2M NaCl, 5 mM β-mercaptoethanol, and 0.01% Triton X-100). Using a peristaltic pump, the high salt buffer was exchanged with low salt buffer (10 mM Tris-HCl [pH 7.5], 1 mM EDTA, 50 mM NaCl, 5 mM β-mercaptoethanol, 0.01% Triton X-100) at roughly 2 ml/min, overnight at 4 °C. In preparation for cryo-EM image collection, the dialysis cassette containing the sample was then placed in a buffer containing 10 mM HEPES-HCl [pH 7.4] and 30 mM KCl, and dialyzed for 48 h at 4 °C. The sample was then incubated at 55 °C for 2 h and centrifuged at 4 °C, 5000 rpm for 5 min in 5415D centrifuge (Eppendorf) to remove aggregates.

For the H1.8 binding nucleosome preparation, recombinant *Xenopus laevis* H1.8 was assembled onto the nucleosome with modifying the previously reported method for linker histone H5 assembly using polyglutamic acid (Stein and Künzler, 1983). Two μM of purified recombinant *X. laevis* H1.8 and 5 ng/μL poly L-glutamic acid (wt 3,000-15,000) (Sigma-Aldrich) were added to 0.15 μM reconstituted nucleosomes in a buffer containing 10 mM HEPES-HCl pH 7.4 and 30 mM KCl. The sample mixture was then incubated at 16 ° for 30 min.

The reconstituted nucleosomes and H1.8 bound nucleosomes were purified and fixed using the GraFix method. Briefly, a sucrose gradient was created using 2.3 ml of a 20 % sucrose solution containing 15 mM HEPES [pH 7.4] and 4 % paraformaldehyde aqueous solution (EM Grade) (Electron Microscopy Sciences) and 2 ml of a 10% sucrose solution containing 15 mM HEPES [pH 7.4]. Reconstituted nucleosome samples were overlaid onto the sucrose gradient and spun at 32000 rpm for 20 h at 4 ° using SW55Ti rotor (Beckman Coulter) in Beckman Optima L80. Fractions containing the fixed samples were collected and dialyzed in 15 mM HEPES [pH 7.4] for 48 h at 4 ° with one fresh exchange of buffer halfway through, using a Tube-O-DIALYZER (100 kDa) (G-Biosciences). Samples were concentrated using an Amicon Ultra 100K centrifugation filter (Millipore) to a final concentration of 195 ng/μl and 247 ng/μl of DNA for the H1.8-binding nucleosome and nucleosome samples, respectively. DNA concentration was calculated using the 260 nm light absorbance.

### Cryo-EM grid preparations and data collection

For the nucleosome purified from *Xenopus* egg extract chromatin, 2 μl of sample was mixed with 0.5 μl of cryoprotectant buffer (10 mM HEPES-KOH [pH 7.4], 30 mM KCl, 1 mM EGTA, 0.3 μg/ml Leupeptin, 0.3 μg/ml Pepstatin, 0.3 μg/ml Chymostatin, 1 mM Sodium Butyrate, 1 mM β-glycerophosphate, 5% trehalose, 0.5% 1,6,-hexanediol) just before loading sample on the cryo-EM grid. 2.5 μl of samples was frozen onto a glow discharged Quantifoil Gold R 1.2/1.3 300 mesh grid (Quantifoil). Samples were frozen under 100% humidity, 20 sec incubation, and 5 sec blotting time using the Vitrobot Mark IV (FEI). Grids were imaged on a Talos Arctica (FEI) installed with a K2 Camera (GATAN) and a field emission gun operating at 200 kV or Titan Krios (FEI) installed with a K2 Camera (GATAN) and a 300 kV field emission gun. All data were collected as 50 frames/movie in super resolution mode. The conditions for the data collection were listed in Table S1 and S2.

### Unbiased Cryo-EM image processing

Both for the interphase oligo fraction and metaphase oligo fraction, data were processed with the identical procedure. Movie frames were motion corrected and dose weighted with a binning factor of 2 using MOTIONCOR2 (Zheng et al, 2017) with RELION3.0 (Scheres, 2012). Particles were picked by blob picker (minimum particle diameter = 50 Å, maximum particle diameter = 300 Å, minimum separation diameters = 100 Å) and extracted (extraction box size = 320 pixels, Fourier crop to 160 pixels) using CryoSPARC v2.15 (Punjani et al., 2017). Extracted particles were applied for 2D classification with 400 classes using CryoSPARC v2.15. Using 2D classification results, particles were split to the lower resolution group and the higher resolution group. The 2D classes whose resolution were higher than 28 Å and effective classes assigned (ECA) value better than 2 were assigned as the higher resolution group. 2D classes containing obvious ice signals, carbon signals, and strong small dots were manually removed from the higher resolution group and reassigned as the lower resolution group. 3D initial models were generated with *ab initio* reconstruction of CryoSPARC v2.15 (10 classes for the higher resolution group, 14 classes for the lower resolution group, maximum resolution = 20 Å, class similarity = 0). 3D classifications were performed using all eighteen 3D classes generated by Ab-initio reconstruction and all picked particles using the heterogeneous refinement of CryoSPARC v2.15 (Refinement box size = 160 voxels). After the 3D classification, particles for each reasonable structure classes were re-extracted with the original pixel size with optimized box sizes for each particle. Nucleosome classes, intelectin classes, actin classes, alpha2-macroglobulin classes were re-extracted with 256, 100, 320, 320 box sizes, respectively. New 3D references were prepared with the re-extracted particles by *ab initio* reconstruction and further refined with homogenous refinement or non-uniform refinement using CryoSPARC v2.15. For the interphase actin and intelectin classes, multiple models were generated by *ab initio* reconstruction (Class similarity = 0) and further “decoy” classifications were performed. Chimera and ChimaraX were used for 3D map visualization (Pettersen et al., 2004).

### Cryo-EM image processing for the high-resolution nucleosome structure determination

All Cryo-EM images of interphase mono-, interphase di-, interphase oligo-, interphase crosslinked after MNase, metaphase mono-, metaphase di-, metaphase oligo-, *in vitro* reconstituted nucleosome, and *in vitro* reconstituted H1.8-bound nucleosome were processed with the same procedure (Fig S4A). Movie frames were motion-corrected and dose-weighted with a binning factor of 2 using MOTIONCOR2 (Zheng et al, 2017) with RELION3.0 or 3.1 (Scheres, 2012). Particles were picked by Topaz v0.2.3 with 500~2000 manually picked nucleosome-like particles as training models (Bepler et al., 2019). Picked particles were extracted using CryoSPARC v2.15 (extraction box size = 200 or 256 pixels) (Punjani et al., 2017). Extracted particles were applied for 2D classification with 200 classes using CryoSPARC v2.15. Using 2D classification result, particles were split to the nucleosome-like groups and the non-nucleosome-like groups. For the interphase di- and metaphase di-data, intelectin 2D classes were removed, and remaining particles were applied for second round of 2D classification, because abundant intelectin-2 particles inhibited the formation of nucleosome-like 2D classes. Four of 3D initial models were generated for both groups with *ab initio* reconstruction of CryoSPARC v2.15 (Class similarity = 0). One nucleosome like model was selected and used as a given model of 3D classification with all four of the “decoy” classes generated from non-nucleosome-like group. After the first round 3D classification, the particles that were assigned to the “decoy” classes were removed, and remained particles were applied for the second round 3D classification with the same setting with the first round. These steps were repeated until 90~95 % of particles were classified as a nucleosome-like class. Repeat times of this step were described in Fig S4A. Using RELION 3.1 beta, CTF refinement, Bayesian polishing, and postprocessing were performed. Local resolution was calculated with the Blocres and Blocfilt in the Bsoft package (Cardone et al., 2013). Chimera and ChimaraX were used for 3D map visualization (Pettersen et al., 2004).

For the 3D classification to analyze linker DNA structure variety shown in Fig 5, S6, and S7, six to eight 3D references were generated with *ab initio* reconstruction of CryoSPARC v2.15 using purified nucleosome-like particles listed in Table S3 (Class similarity = 0.9). 3D classifications was performed using all newly generated 3D references and purified particles using the heterogeneous refinement of CryoSPARC v2.15.

For the atomic model building, initial atomic models of linker DNA containing nucleosomes were built by combining linker DNA moiety of gH1.0 binding nucleosome (PDB ID: 5NL0) and nucleosome moiety of ScFv binding nucleosome (PDB ID: 6dzt) and by replacing amino acids sequence to that of *Xenopus laevis* core histones using Coot (Bednar et al., 2017; Emsley and Cowtan, 2004; Zhou et al., 2019). The initial atomic model was docked onto the cryo-EM map postprocessed by RELION 3.1 beta and refined using Coot and PHENIX (Liebschner et al., 2019).

### 3D Variability Analysis (3DVA)

For the 3D structure dynamics analysis of nucleosome containing linker DNA shown in Fig 4A, the purified nucleosome particles used for 3D classification analysis (listed in Table S3) were used for the 3DVA in Cryo-SPARC v2.15 (Punjani and Fleet, 2020). For all samples, the number of 3DVA components was set to 2. For the interphase nucleosomes crosslinked after MNase, filter resolution was set to 12 Å, whereas for other samples, filter resolution was set to 8 Å. All output 3D maps were loaded with Chimera (Pettersen et al., 2004) and movies were created from the maps.

For the 3DVA of NCPs shown in Fig 4B, C, linker DNA densities of each nucleosome were removed from original particles listed in Table S3 using particle subtraction in cryoSPARC v2.15.

For GUB NCPs, the new initial 3D map was generated with linker DNA subtracted particles by *ab initio* reconstruction of cryoSPARC v2.15. The 3D map was further refined with homogeneous refinement of cryoSPARC v2.15. Particle stack and mask generated by NU-refinement were used for 3DVA of cryoSPARC v2.15 (number of modes = 2, Filter resolution = 8Å).

For chromosomal NCPs, 100,000 each particle was selected from interphase or metaphase nucleosome particles whose linker DNAs had been subtracted. Particles were mixed, and the new initial 3D map was generated by ab initio reconstruction of cryoSPARC v2.15. Particle stack and mask generated by homogeneous refinement were used for 3DVA of cryoSPARC v2.15 (number of modes = 4, Filter resolution = 8Å).

To calculate the surface hydrophobicity and electrostatic potential and global R.M.S.D., an atomic model was generated for each NCP structural variant. Using PHENIX package, the atomic model of interphase oligo nucleosome was docked on to the 3D maps of NCP structural variants generated by 3DVA. Docked atomic models were refined with rigid body refinement in PHENIX. Global R.M.S.D. of Cα and P atoms were calculated by PyMol. To show the surface hydrophobicity (Fig. 7C), hydropathy indexes (Kyte and Doolittle, 1982) were assigned to each amino acid residue and mapped on the original EM maps by Chimera. To show the electrostatic potential, surface electrostatic potentials were calculated by APBS and PDB2PQR (Baker et al., 2001; Dolinsky et al., 2004) using refined atomic models for each NCP structural variant, and the colored surface electrostatic potentials were drawn by Chimera.

### *In silico* mixing 3D classification

For the *in silico* mixing 3D classification for analyzing linker DNA structure variation shown in Fig 5 and S6, five different linker DNA angle 3D maps were selected as the fixed templates for all samples (“closed” : metaphase oligo class A (Fig 5B), “601-like” : metaphase oligo class D (Fig 5B), “open” : interphase oligo class E (Fig 5A), “wide open” : interphase crosslink after MNase class E (Fig 5E), and “flexible DNA” : metaphase oligo class F (Fig 5B)). The same number of particles (mono; 13,000 particles, di : 6,000 particles, oligo : 25,000 particles) were randomly selected from the purified particle set (listed in Table S3) of metaphase nucleosome, interphase nucleosome, and interphase nucleosome crosslinked after MNase. After merging these particles, 3D classification was performed five times using all five 3D maps and merged particle set using heterogeneous refinement of CryoSPARC v2.15 (refinement box size =200, initial resolution = 32 Å). The particles assigned to each class were counted and plotted.

For the *in silico* mixing 3D classification shown in Fig. 7C, three 3D maps (frame 4, 11, and 18) for each component were selected from 3DVA of GUB NCP, which had generated 21 motion frames per each component. In the 3DVA, largest structural differences are found between motion frames 1 and 21, and they are structurally most deviated from the averaged GUB NCP structure, while the structure of the motion frame 11 is most similar to that of the averaged GUB NCP. We chose frames 4 and 18, instead of frames 1-3 and 19-21, for reference maps because frames 1 and 21 contain many noisy densities (e.g., gaps in helices and densities outside of the NCP). As a 3D reference of canonical nucleosome structure in chromosome, a refined 3D map generated from 100,000 each of linker DNA subtracted interphase oligo nucleosome, metaphase oligo nucleosome particle was used. As decoy maps, 4 noise classes shown in Fig. S4 were used. As input particles for *in silico* mixing 3D classification, 25,000 each of linker DNA subtracted interphase oligo nucleosome, metaphase oligo nucleosome, interphase nucleosome crosslinked after MNase, and GUB nucleosome particles were randomly selected and mixed. Using these 3D reference maps and particles, 3D classification was performed d five times using heterogeneous refinement of CryoSPARC v2.15 (refinement box size =200, initial resolution = 20 Å, number of initial random assignment iterations = 0). To confirm the accuracy of 3D classification, 3D maps were newly generated with output particles of *in silico* mixing 3D classification by *ab initio* reconstruction and NU-refinement. Newly generated structures were compared to the input maps of the *in silico* mixing 3D classification, and the parameters of the *in silico* mixing 3D classification were optimized for newly generated structures to preserve the feature of input maps (data available upon request). For each 3DVA component, the *in silico* mixing 3D classification was performed five times. The particles assigned to each class were counted and plotted.

For the *in silico* mixing 3D classification shown in Fig. 7D, three 3D maps (frame 4, 11, and 18) of each component generated by 3DVA of chromosomal NCP were selected. The 3DVA of chromosomal NCP (using metaphase and interphase oligo nucleosome particles) generated 21 motion frames per each component. Frames 11 of each component are most similar to the canonical nucleosome structure in chromosome, while we chose frames 4 and 18 of components 1-4 to represent NCP structural variants, as the rationale explained above for the GUB nucleosome *in silico* mixing 3D classification. As decoy maps, 4 noise classes shown in Fig. S4 were used. As input particles for *in silico* mixing 3D classification, 100,000 each of linker DNA subtracted interphase oligo nucleosome and metaphase oligo nucleosome particles were randomly selected and mixed. Using these 3D reference maps and particles, 3D classification was performed five times using heterogeneous refinement of CryoSPARC v2.15 (refinement box size =200, initial resolution = 13 Å, number of initial random assignment iterations = 0). To confirm the accuracy of 3D classification, 3D maps were newly generated with output particle sets of *in silico* mixing 3D classification by *ab initio* reconstruction and NU-refinement. Newly generated structures were compared to the input maps of the *in silico* mixing 3D classification, and the parameters of the *in silico* mixing 3D classification were optimized for newly generated structures to preserve the feature of input maps (data available upon request). For each 3DVA component, the *in silico* mixing 3D classification was performed five times. The particles assigned to each class were counted and plotted.

### Reconstruction of “closed” and “moderately open” linker DNA nucleosome maps (Fig. 6G)

Using the interphase nucleosome particles shown in Table S3, twenty classes of 3D maps were generated by 3DVA of cryoSPARC v2.15 (cluster mode, number of modes = 2, Filter resolution = 8Å, number of frames = 20). To generate the “closed” and “open” linker DNA nucleosome maps at the comparable resolution, five “closed” linker DNA classes (20,738 particles) and five “moderately open” linker DNA classes (19,120 particles) were selected, and initial 3D maps were generated by *ab initio* reconstruction of cryoSPARC v2.15. Initial 3D models were further refined with homogeneous refinement in cryoSPARC v2.15.

### Cryo-EM image processing for the H1.8-bound nucleosome structure determination

41,416 particles assigned to the class that has the extra density of the metaphase oligo fraction were applied for homogeneous refinement (Fig S7A). Refined particles were further purified with the heterogeneous refinement using an H1-bound class and an H1-unbound class as decoys. The purified particles were further refined and split to four of 3D classes using the *ab initio* reconstruction of (4 classes, class similarity = 0.9) and heterogeneous refinement of CryoSPARC v2.15. Two classes that had reasonable extra density were selected and refined with homogeneous refinement. The final resolution was determined as 4.42Å with the gold stand FSC threshold (FSC = 0.143).

For the atomic model building, H1 segment of the gH1.0-bound nucleosome structure (PDB ID: 5NL0) was replaced with the H1.8 structure model built using SWISS-MODEL: homology modeling tool (Waterhouse et al., 2018). The initial atomic model was docked onto the cryo-EM map locally filtered by Bsoft (Blocres and Blocfilt) and refined using Coot and PHENIX (Cardone et al., 2013; Emsley and Cowtan, 2004; Liebschner et al., 2019). Chimera and ChimaraX were used for 3D map visualization (Pettersen et al., 2004).

---

**Figure S1.**
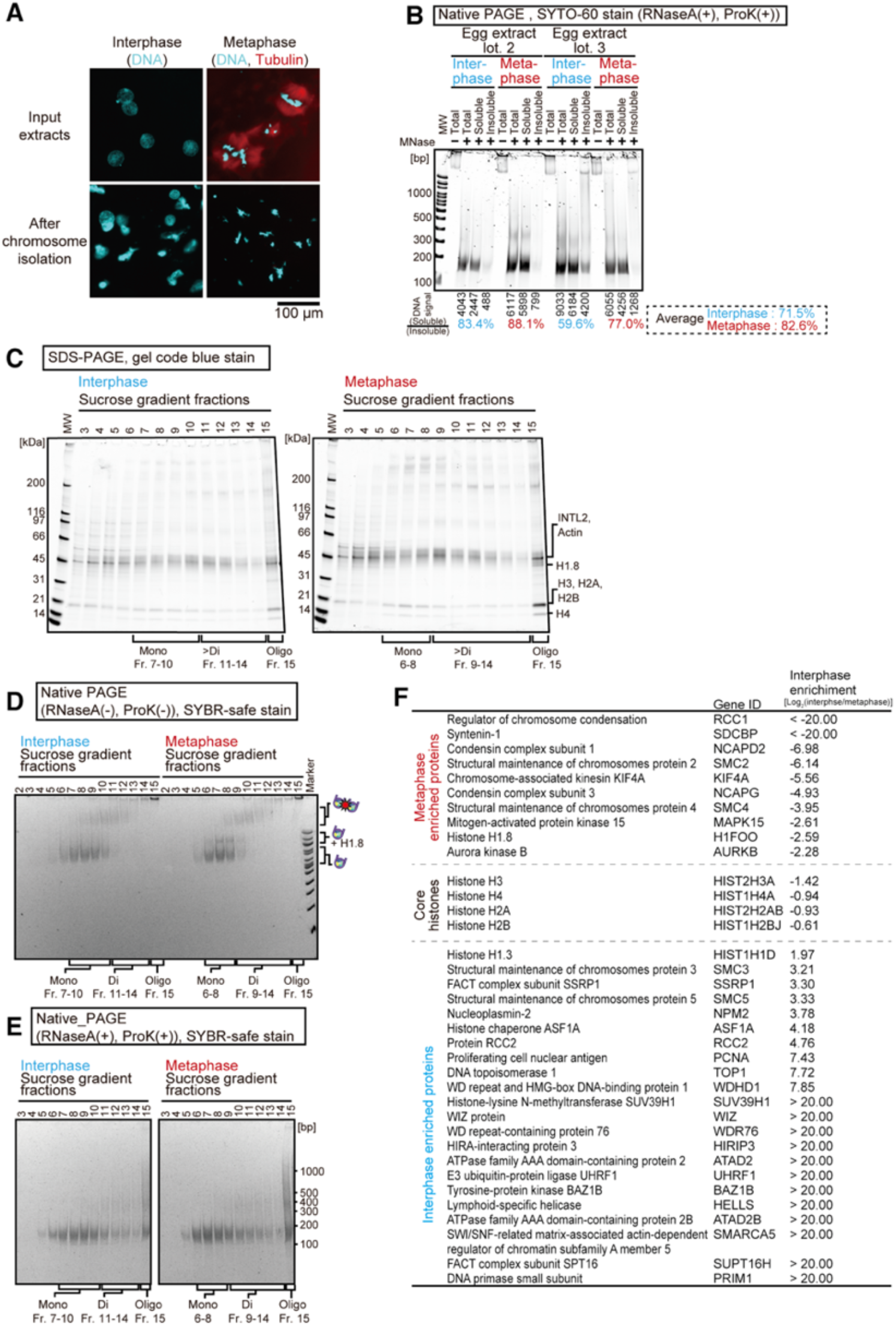
Isolation of nucleosomes from interphase and metaphase chromosomes. Related to Figure 1. **A**, Top panels, interphase nuclei and metaphase sperm chromosomes assembled in the *Xenopus* egg extract. Bottom panels, interphase nuclei and metaphase chromosomes after isolation. Fluorescence images of Hoechst 33258-stained DNA and Alexa594-labeled tubulin are shown. **B**, Quantification of the chromatin solubility after MNase. Chromosomes were isolated from 800 μL of two independent lots of *Xenopus* egg extracts with our chromatin isolation protocol. 20 μL of MNase treated chromatin was mixed with 80 μL of MNase stop buffer and spun at 13,894 rpm (16,000 rcf) at 4 °C for 30 min using a SX241.5 rotor in Allegron X-30R (Beckman Coulter). Insoluble pellets were resuspended with 20 μL of buffer SC and 80 μL MNase stop buffer. 5 μL each of soluble and insoluble fractions were mixed with 1 μL of 10 mg/ml RNaseA (Thermo Scientific) and incubated at 55 °C for 30 min. RNaseA treated samples were then mixed with 1 μL of 19 mg/ml Proteinase K solution (Roche) and incubated at 55 °C overnight. Samples were loaded onto a 0.5x TBE 6 % native PAGE. DNA was stained by SYTO60. Signal intensities were measured with ImageJ. **C**, SDS-PAGE analysis of the nucleosome fractions isolated from *Xenopus* egg extract sperm chromatin. 15μl of each nucleosome fractions were reverse-crosslinked prior to loading onto a 4-20 % gradient gel. Proteins were stained by gel code blue. Fraction 1 and fraction 15 represent the top and bottom of the sucrose gradient, respectively. For interphase samples, fractions 7 to 10, 11 to 14, and 15 represent ‘mono’, ‘di’, and ‘oligo’ fractions, respectively. For metaphase samples, fractions 7 to 10, 11 to 14, and 15 represent ‘mono’, ‘di-’, and ‘oligo’ fractions, respectively **D**, Native PAGE analysis of the nucleosome fractions isolated from sperm chromosomes assembled in *Xenopus* egg extracts. 15 μl of each nucleosome fractions was loaded onto a 0.5x TBE 6% native PAGE. DNA was stained by SYBR-safe. **E**, Native PAGE analysis of the nucleosomal DNA isolated from sperm chromosomes assembled in *Xenopus* egg extracts. 15 μl of each nucleosome fractions was deproteinized and reverse-crosslinked prior to loading onto a 0.5x TBE 6% native PAGE. DNA was stained by SYBR-safe. **F,** Cell cycle specific chromatin-associated proteins detected by MS of the interphase or metaphase oligo fractions.

**Figure S2.**
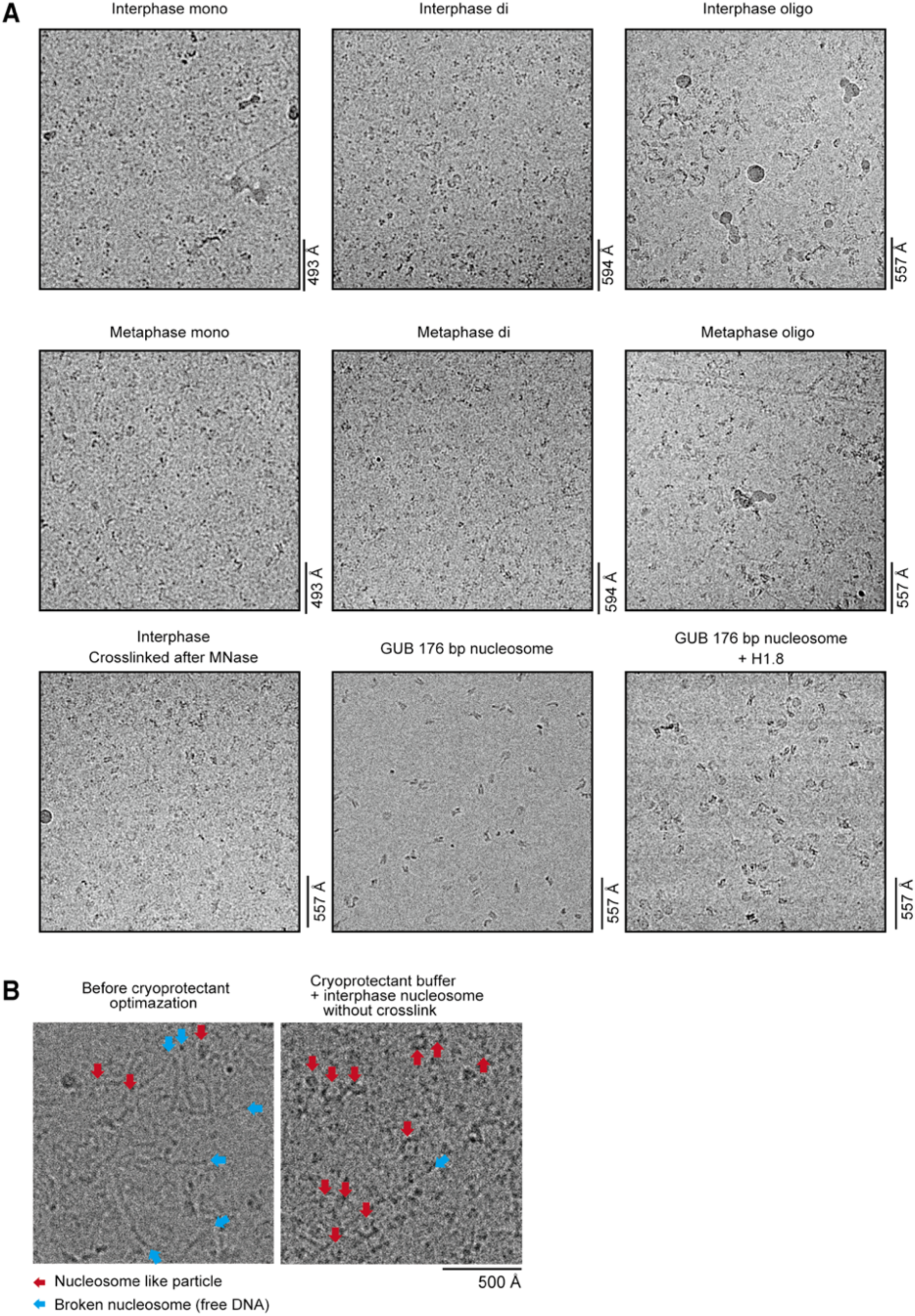
Raw micrographs and picked particles for the unbiased reconstruction. Related to Figures 2 and 3. **A**, Raw micrographs used in this study. Each micrograph represents a quarter of the original image. **B,** Raw micrographs with interphase nucleosomes without crosslinking either in the presence or absence of optimized cryoprotectant buffer. Broken DNA fragments can be observed in samples that were frozen in the absence of cryoprotectant buffer.

**Figure S3.**
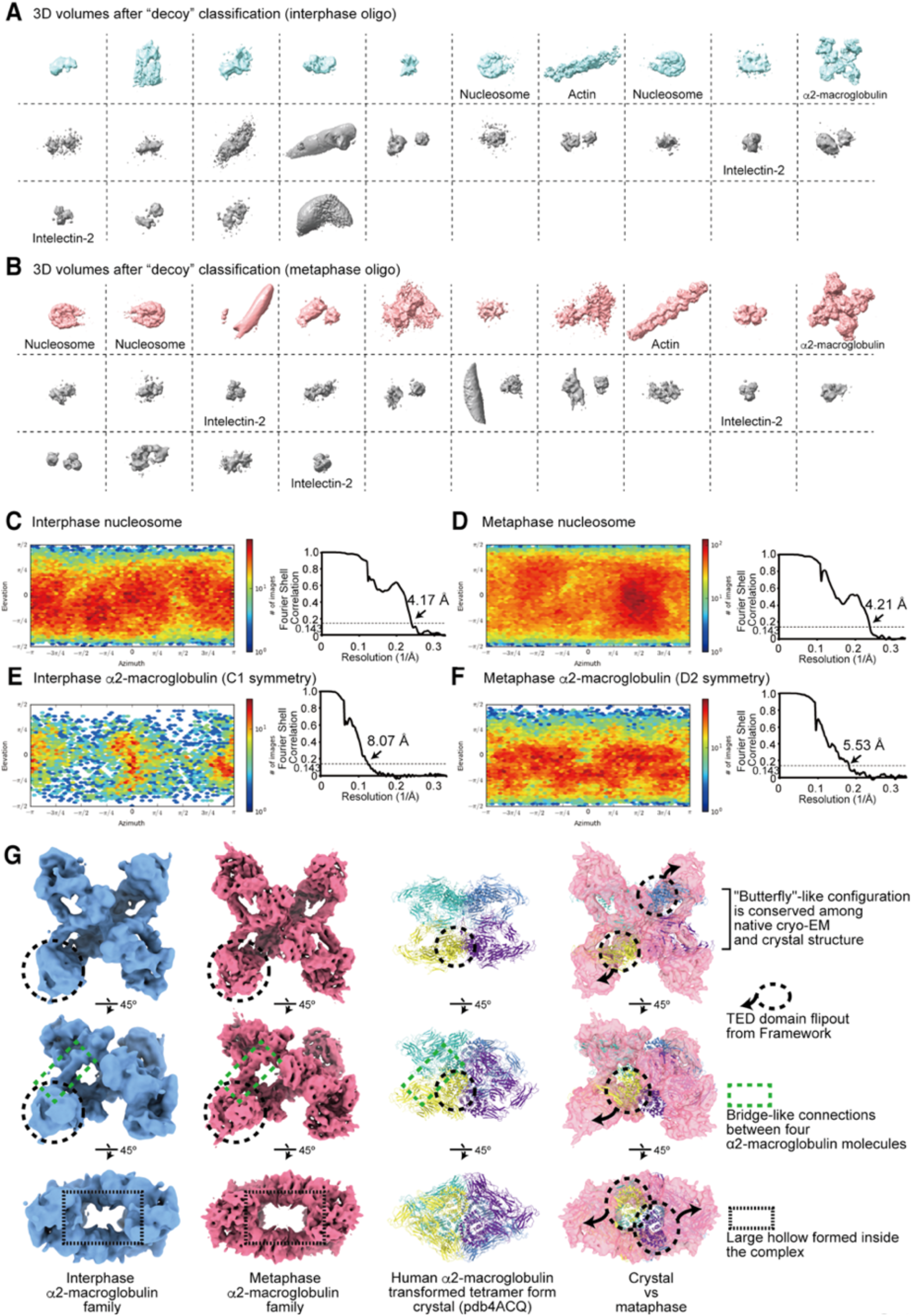
Unbiased reconstruction of cryo-EM structures. Related to Figure 2. **A, B**, 3D classes obtained by the decoy 3D classification of interphase (A) and metaphase (B) particles. **C, D,** Angular distribution plots and Fourier shell correlations of the nucleosome structure obtained from the interphase oligo fraction (C) and the metaphase oligo fraction (D). **E, F,** Angular distribution plot and Fourier shell correlation of the alpha2-macroglobulin family protein structure obtained from interphase (E) and metaphase (F) oligo fractions **G,** Structural comparison between the cryo-EM structure of *X.laevis* native alpha2-macroglobulin family protein determined in this study and the crystal structure of human recombinant alpha2-macroglobulin (PDB ID 4ACQ). While the “butterfly” like-subdomain structure was conserved between the cryo-EM of the native *Xenopus* complex and the crystal structure of the purified human proteins, the major change in the overall structures is observed at the TED domain, which appears to flip out in the cryo-EM structure from the crystal structure. Since the cryo-EM structured was determined on alpha2-macroglobulin from the cytoplasm fraction of Xenopus eggs, the structure is expected to undergo a major conversion when it becomes processed into the active form in the extracellular space.

**Figure S4.**
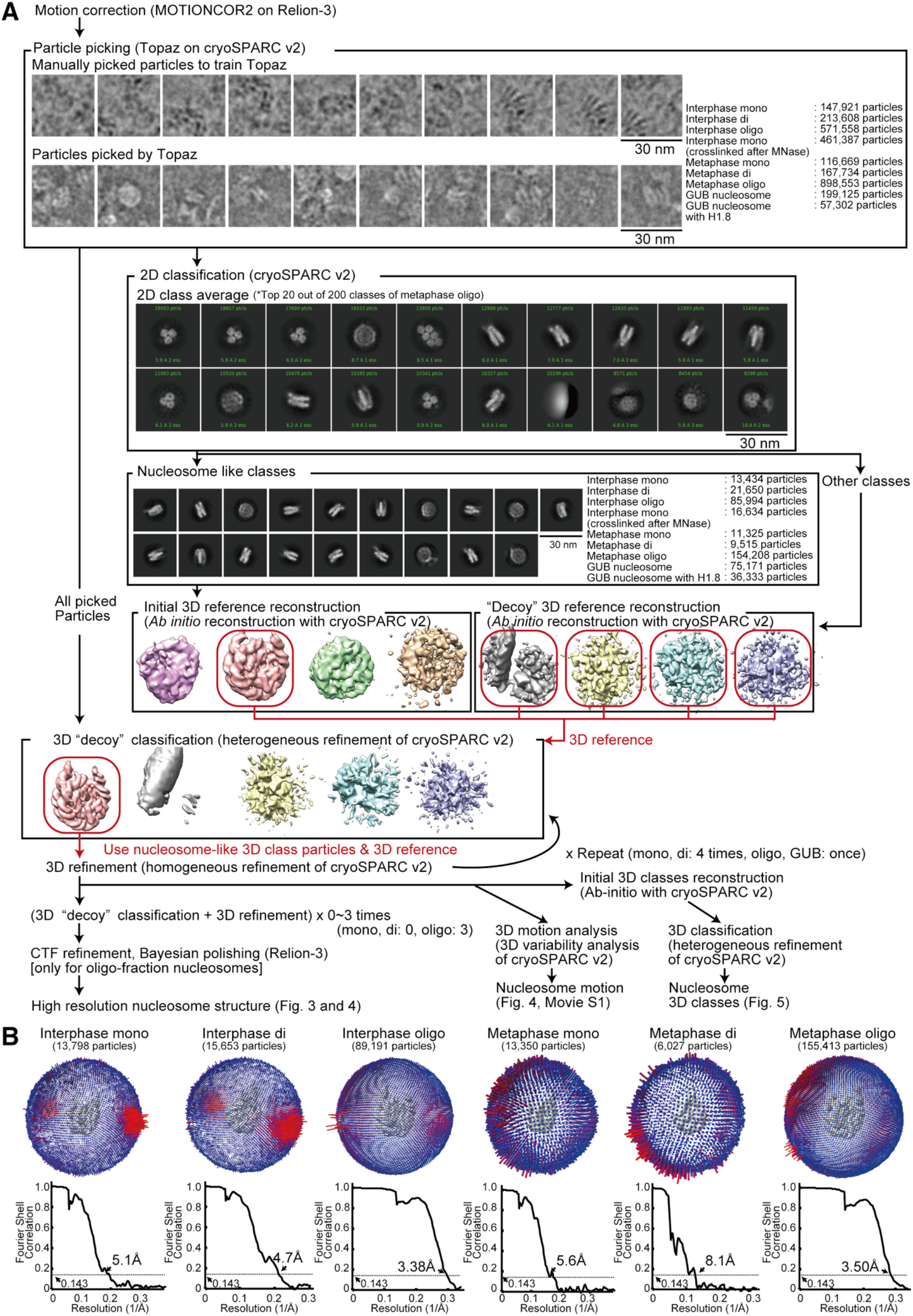
Pipeline of the high-resolution nucleosome structure analysis. Related to Figure 3. **A,** The high-resolution nucleosome structure determination pipeline. Picked particle images, 2D classes, 3D structures from the metaphase oligo fraction are shown. **B,** Angular distribution plots and Fourier shell correlations of the reconstructed nucleosome structures.

**Figure S5.**
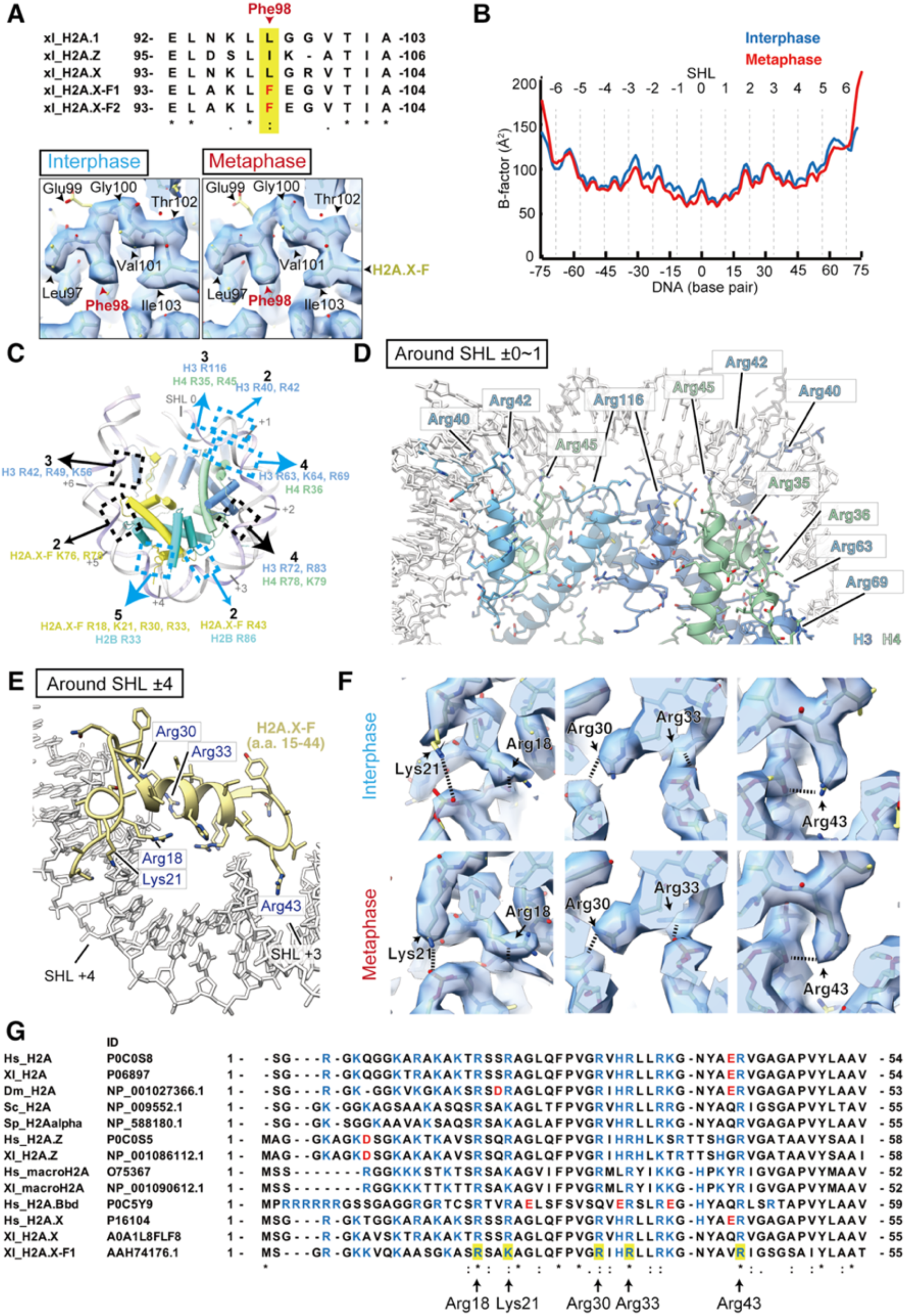
DNA positioning at SHL ±0~1 and SHL ±3.5~4.5 are well ordered in both interphase and metaphase nucleosomes. Related to Figure 3. **A**, Identification of Phe98 of H2A.X-F1/2, the dominant H2A variant in eggs on solved cryo-EM density maps. Amino acid sequence alignment of H2A variants, and cryo-EM densities of the regions of interest in interphase and metaphase nucleosomes. **B,** B-factors of the DNA in the interphase and metaphase nucleosomes from oligo fractions. B-factors of nucleotides were calculated using PHENIX real-space refinement (Liebschner et al., 2019). **The** B-factor of each DNA base pair was calculated by averaging the B-factors of two nucleotides forming the base pairing. **C**, Number and identity of basic amino acid residues that interact with DNA. Hydrogen atoms were added to the atomic coordinate of metaphase nucleosome and the chemical interaction sites were extracted using RING 2.0 webserver (Piovesan et al., 2016). The basic amino acid residues with EM density in close contact (< 3.0 Å) with DNA were mapped on the nucleosome 3D structure. Blue arrows indicate potential interactions contributing to the DNA positioning stabilization at SHL ±0~1 and SHL ±3.5~4.5. **D**, The DNA-histone interactions in the metaphase nucleosome structure at SHL ±0~1. **E,** Interactions between H2A.X-F and DNA at SHL ±3.5~4.5 in metaphase nucleosomes. **F,** EM maps around the DNA-histone interaction sites at SHL ±3.5~4.5. **G**, Amino acid sequence alignment of the H2A N-terminal region. Residues forming hydrogen bonds with DNA at SHL ±0~1 are highlighted. Hs, *Homo sapiens*; Xl, *Xenopus laevis*; Sc, *Saccharomyces cerevisiae*; Sp, *Schizosaccharomyces pombe*.

**Figure S6.**
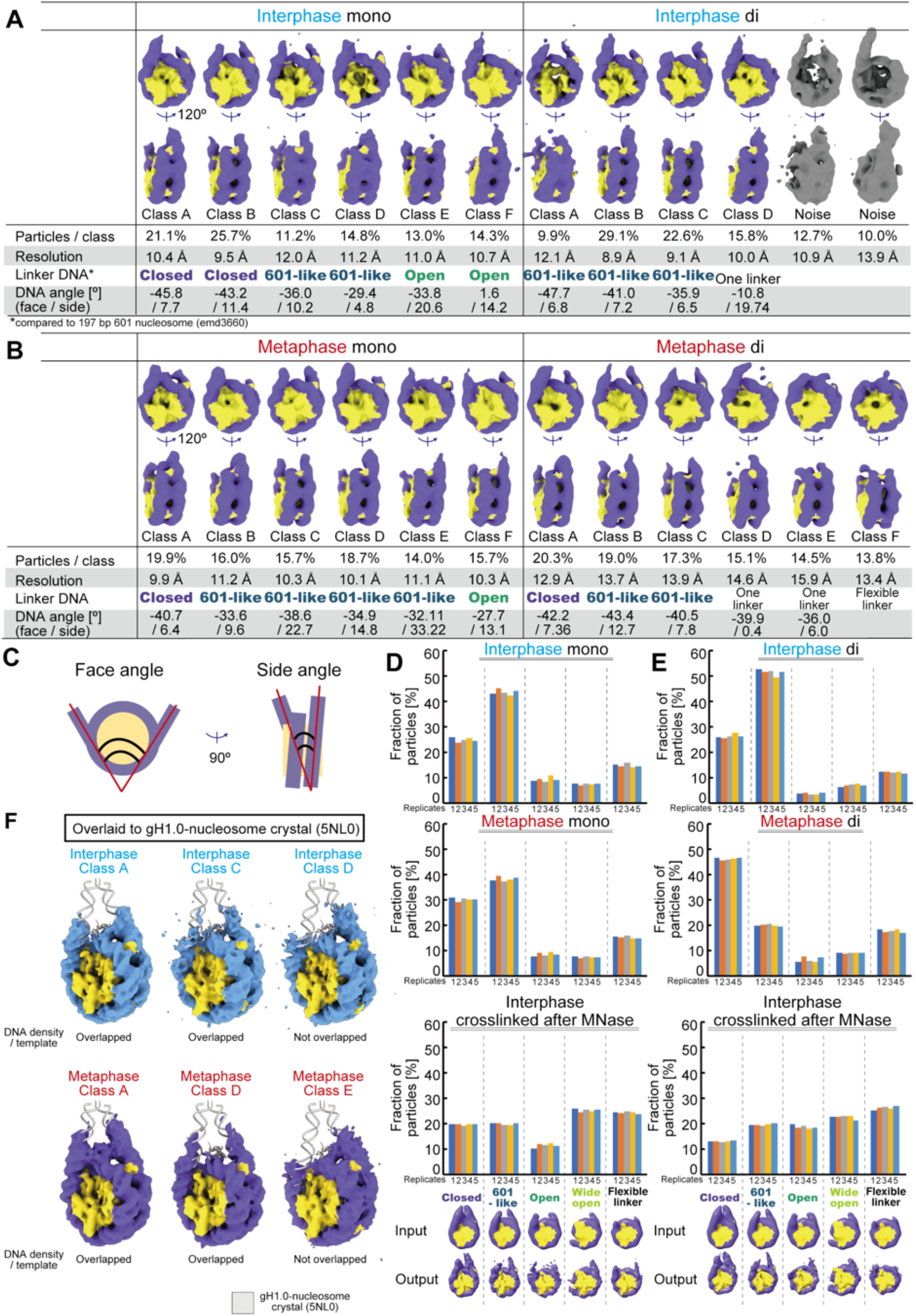
Linker DNA angles of a majority of interphase and metaphase nucleosomes are closed. Related to Figure 5. **A**, **B,** 3D structure classes of the nucleosome from the mono and di fractions of interphase (A) and metaphase (B) nucleosomes. **C**, Linker DNA angle definitions. **D, E,** 3D classification reproducibility analysis with fixed 3D templates for the mono (D) and di (E) fractions of nucleosomes. 3D classification calculations were performed with five different linker DNA angle 3D maps low-pass filtered to 32 Å. The relative fractions assigned to each class of the five independent 3D classification analyses are shown. 3D map images at the bottom indicate the input and output of the 3D classifications. **F**, Structural comparison between nucleosome structure classes in chromosomes and the crystal structure of gH1.0-bound nucleosome (PDB ID: 5NL0). Cryo-EM map of classes A, C, and D of the interphase nucleosome (oligo fraction), and classes A, C, and D of the metaphase nucleosome (oligo fraction) were overlaid onto the atomic model of the gH1.0 bound nucleosome structure (shown in gray). The blue map indicates the DNA of the interphase nucleosome and the purple map indicates DNA of the metaphase nucleosome.

**Figure S7.**
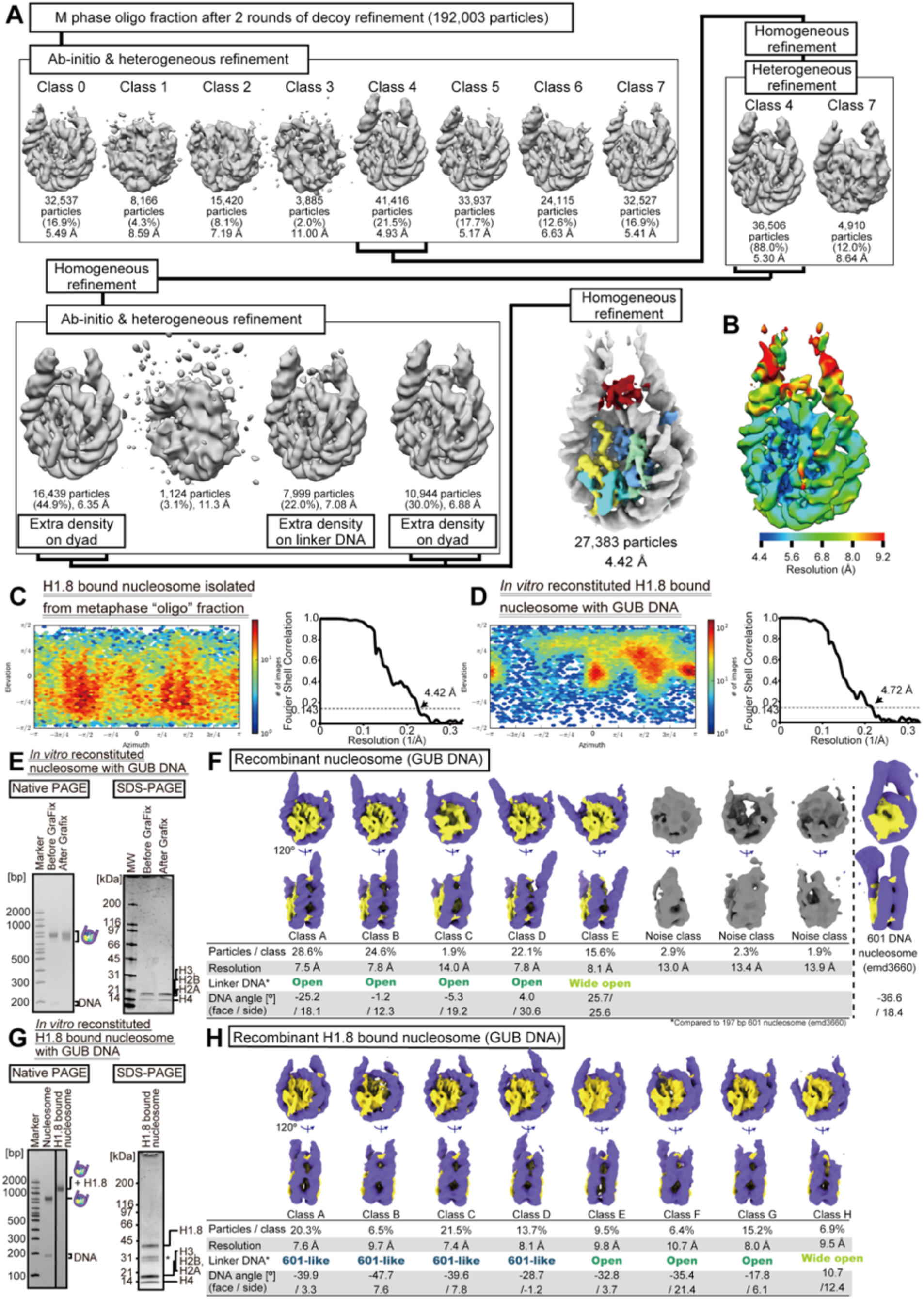
Cryo-EM analyses of H1.8-bound nucleosomes. Related to Figure 6. **A,** Pipeline for isolating the class that has an extra density on nucleosome dyad. **B,** Local resolution of the H1.8-bound nucleosome, calculated with BLOCRES and BLOCFILT in the BSOFT package. **C,** Angular distribution plot and Fourier shell correlation of the H1.8-bound nucleosomes isolated from chromosomes in metaphase *Xenopus* egg extract. **D,** Angular distribution plot and Fourier shell correlation of nucleosomes reconstituted with recombinant core histones, GUB nucleosome-positioning sequence, and recombinant H1.8 *in vitro*. **E,** SDS-PAGE and native PAGE of the purified *in vitro* reconstituted nucleosome. **F,** 3D structure classes of the *in vitro* reconstituted nucleosome. **G,** SDS-PAGE and native PAGE of the purified *in vitro* reconstituted H1.8-bound nucleosome. **H,** 3D structure classes of the *in vitro* reconstituted H1.8-bound nucleosome.

**TableS1.**
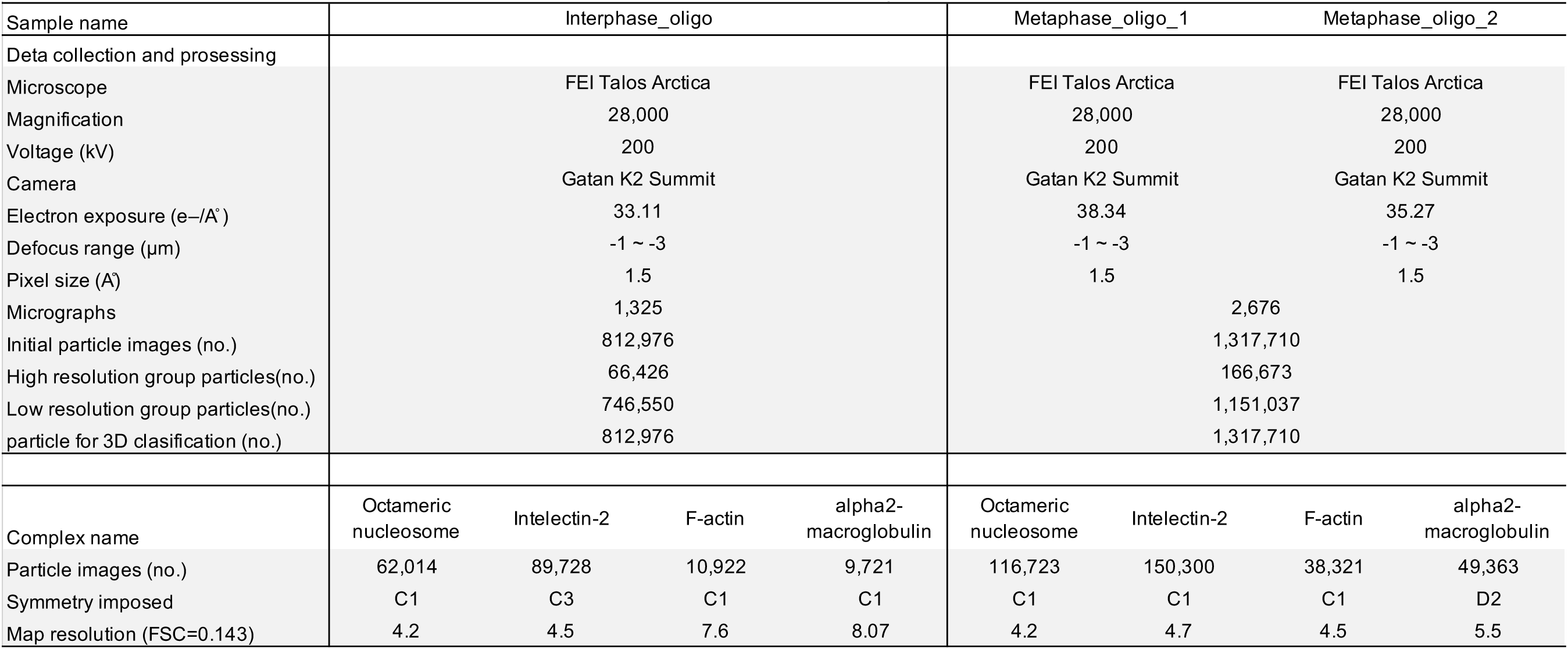
Data collection, model refinement and validation for “unbiased” reconstruction. Related to Figure 2

**TableS2.**
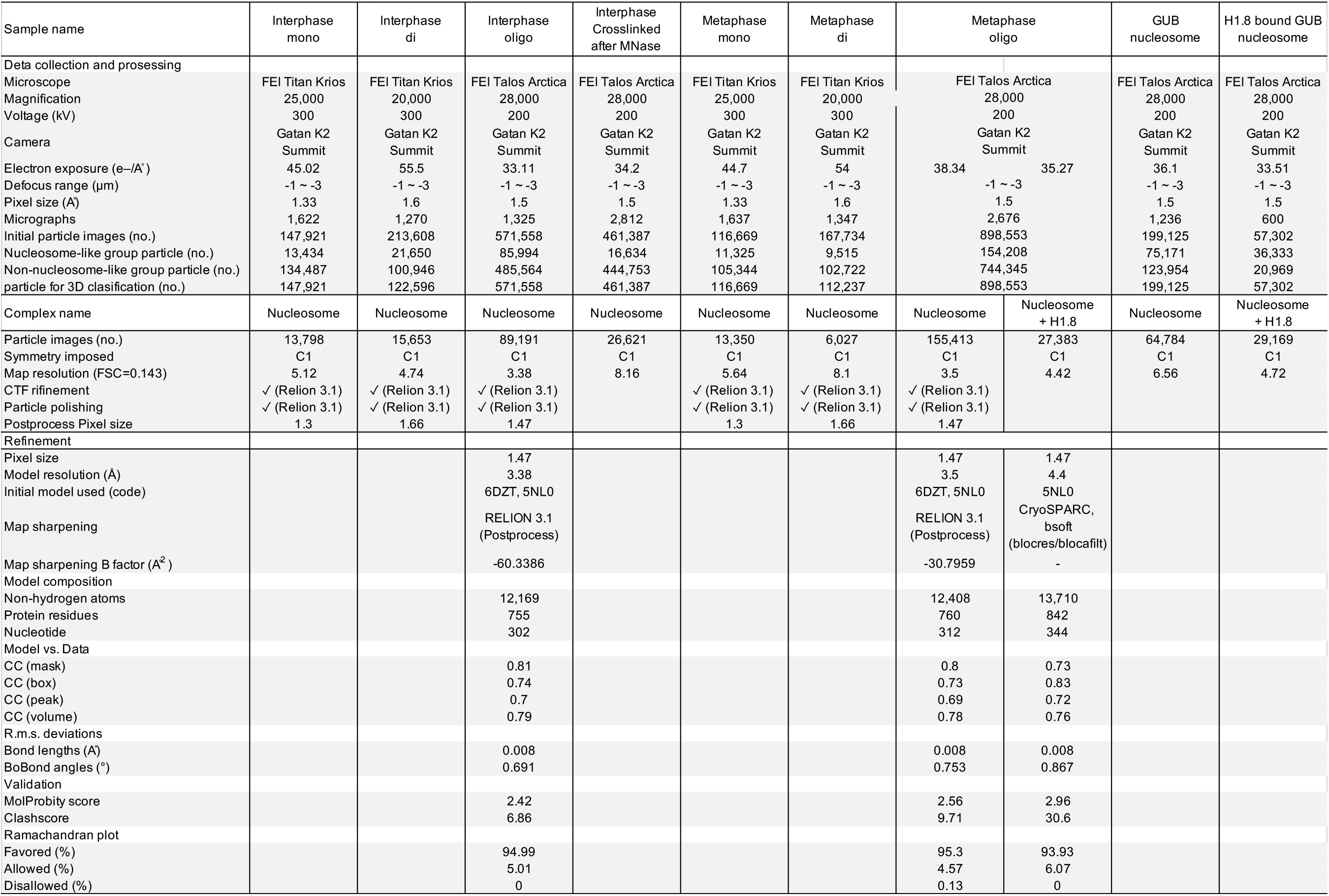
Data collection, model refinement and validation for “high resolution” structure analysis. Related to Figure 3

**TableS3.** Number of perticles and classes of 3D classification. Related to Figure 5 and 6

|  | Particles | Classes |
| --- | --- | --- |
| Interphase mono | 13,798 | 6 |
| Interphase di | 15,653 | 6 |
| Interphase oligo | 106,453 | 8 |
| Interphase nucleosome crosslinked after MNase | 26,621 | 8 |
| Metaphase mono | 13,350 | 6 |
| Metaphase di | 6,027 | 6 |
| Metaphase oligo | 192,003 | 8 |
| GUB nucleosome | 64,784 | 8 |
| GUB nucleosome with H1.8 | 29,169 | 8 |

